# Loss of competitive strength in European conifer species under climate change

**DOI:** 10.64898/2026.02.13.705703

**Authors:** Marc Grünig, Werner Rammer, Martin Baumann, Katharina Albrich, Frédéric André, Andrey L. D. Augustynczik, Friedrich Bohn, Meike Bouwman, Harald Bugmann, Alessio Collalti, Irina Cristal, Daniela Dalmonech, Miquel De Caceres, Francois De Coligny, Laura Dobor, Christina Dollinger, David I. Forrester, Jordi Garcia-Gonzalo, José Ramón González-Olabarria, Ulrike Hiltner, Tomáš Hlásny, Juha Honkaniemi, Nica Huber, Andreas Huth, Mathieu Jonard, Anna Maria Jönsson, Fredrik Lagergren, Marco Mina, Frits Mohren, Christine Moos, Xavier Morin, Bart Muys, Mats Nieberg, Mikko Peltoniemi, Christopher PO Reyer, Ilié Storms, Dominik Thom, Maude Toigo, Rupert Seidl

## Abstract

Climate change is expected to alter species assemblages by affecting the outcome of competition between species. Investigating processes of competition remains challenging particularly in tree communities, as they unfold over extensive spatio-temporal scales. Here, we developed a deep-learning approach to leverage a novel database of 135 million simulated local-scale tree responses to climate across continental Europe to investigate changes in the competitiveness of nine major tree species under different scenarios of climate change. Specifically, we trained a Deep Neural Network on local process model projections to investigate climate change effects on indicators of competitive strength and species dominance. We found decreasing competitive strength for all investigated evergreen coniferous species across their distribution, while major deciduous broadleaved species such as *Quercus robur* and *Fagus sylvatica* increased in competitiveness. Changes in tree species competition with climate differed locally, but most investigated species lost competitive strength at their warm range edges. As a consequence of these changes, up to 19% of Europe’s forests could experience a change in the dominant tree species until the end of the 21^st^ century. Our results suggest a profound climate-induced reassembly of Europe’s forests and identify areas that may require specific attention in forest policy and management.

## Introduction

Competition for limited resources is a fundamental process driving the composition and structure of vegetation around the globe ^1–3^. Competition is a mutual process between co-occurring species striving for the same resources, and its outcome is determined by the competitiveness and sensitivity to competition of the respective species ^4,5^. In forested ecosystems, competition between tree species is driven by traits that determine the potential to overgrow others (i.e., height growth potential) ^6^, the ability to shade out competitors (i.e., high leaf area index), tree longevity (i.e., the ability to outlive competitors), and the ability to acquire water and nutrients more efficiently than neighbouring trees (e.g., different rooting strategies) ^7,8^. Although facilitation may occur between trees, competition remains the dominant process of interaction between trees in forest ecosystems ^9,10^. Under stable environmental conditions, highly competitive tree species can maintain their dominance over extended periods of time and cover extensive areas. European beech (*Fagus sylvatica* L.), for instance, is able to overgrow most other tree species and is highly shade tolerant; thus it dominates natural forest composition in large parts of temperate Europe ^11^.

Changing environmental conditions due to climate change can have profound impacts on the competitiveness of tree species, as they shift the availability and distribution of critical resources in space and time ^12^. As a consequence, species could lose competitiveness or experience higher sensitivity to competition, resulting in reduced competitive strength relative to other species and ultimately leading to a change in species assemblages ^13,14^. Therefore, the loss of competitive strength of individual species can serve as an early warning indicator for climate-induced shifts in tree species composition ^15^. Such shifts could affect the resource use efficiency and carbon uptake of forest ecosystems, as well as other important ecosystem services ^16^. Losses in competitive strength could also be significant for biodiversity, e.g., when species associated with a particular tree species lose their resource base due to the decline of their host species ^17,18^. Consequently, a quantification of potential changes in the competitive strength of tree species is highly relevant for predicting ecosystem responses to ongoing environmental changes.

Assessing changes in the competitive strength of trees is challenging, as competition occurs over extended periods of time. Furthermore, competitive strength can vary throughout the range and life stages of a species, limiting the utility of short-term, local assessments and requiring the consideration of large spatio-temporal extents. Process-based simulation models relying on fundamental principles of forest dynamics are prime tools to overcome these challenges ^19^. They can be applied to study the outcomes of competition over timeframes of decades to centuries consistently across a range of locations, while explicitly accounting for the effects of changing climatic conditions. A number of process-based forest models have been developed over the past decades, designed to capture local ecosystem characteristics and parametrized and tested with local data. These models thus represent our best available quantitative understanding of how forests in a given region might respond to climate change ^20^. Yet, this treasure trove of local climate change responses has not been synthesized across locations and models, and thus remains underutilized in large-scale assessments and policy considerations. Recent breakthroughs in artificial intelligence (AI) present new opportunities to learn from the nuanced climate change responses observed in local simulation data, e.g. by conducting AI-based model synthesis ^21,22^.

Here, we harnessed a novel database of harmonized local forest simulations across Europe containing 135 million simulation-years from 17 process-based forest models ^23^ to train a Deep Neural Network (DNN) of forest dynamics. Subsequently, we used this DNN meta-model to investigate how climate change affects competitive strength. Specifically, we (i) examined the change in competitive strength for nine widely distributed European tree species (i.e., European beech (*Fagus sylvatica* L.), Scots pine (*Pinus sylvestris* L.), silver fir (*Abies alba* Mill.), Norway spruce (*Picea abies* (L.) H. Karst.), European larch (*Larix decidua* Mill.), silver birch (*Betula pendula* Roth), Aleppo pine (*Pinus halepensis* Mill.), pedunculate oak (*Quercus robur* L.), and holm oak (*Quercus ilex* L.)) within their current range limits at the level of individual species and (ii) highlighted in which geographical regions and sections of their current niche (i.e., warm vs. cold edges of the distribution) species gain or lose competitive strength under climate change. For these analyses we used height growth and leaf area index (LAI) as indicators of competitive strength, and combined them to a competitive strength index (CSI). Subsequently, we analyzed (iii) whether the potential loss in competitive strength at species level also results in a change in species dominance at population level, i.e. if changes in species-level competitive strength also result in changes in tree species composition. We furthermore (iv) identified hotspots of shifting species dominance, pinpointing areas in which assemblages are likely to change and which require special attention in forest policy and management.

## Results

### Change in competitive strength of major European tree species

We found that six out of nine studied species, including all investigated evergreen species, experienced a decline in competitive strength under climate change (Fig. 1). Under severe climate change (scenario RCP8.5, period 2071-2100), the competitive strength (CSI) of Aleppo pine (−22.6%), holm oak (−5.3%), and Norway spruce (−5.3%) decreased across their current distribution ranges, relative to values under current climate (1981-2010). In addition, also silver birch (−4.8%), silver fir (−4.3%) and Scots pine (−1.5%) had negative CSI values. In contrast, pedunculate oak (+7.1%), European beech (+2.6%) and European larch (+1.6%) increased their competitive strength under climate change. On average, broadleaved species responded slightly positive to climate change (+0.1%), while the competitive strength of coniferous species decreased (−3.2%). While patterns remained robust also under strong climate change (RCP4.5), only the two Mediterranean species Aleppo pine (−7.7%) and holm oak (−2.6%) decreased in competitive strength under moderate climate change (RCP2.6). Norway spruce and silver birch, which both showed decreasing competitive strength under RCP8.5, benefitted under RCP2.6 (+3.4%; +0.6%). Analyzing the two components of CSI, we found that LAI responded more negatively than height growth across most species (Fig. S1 & S2).

**Figure 1.**
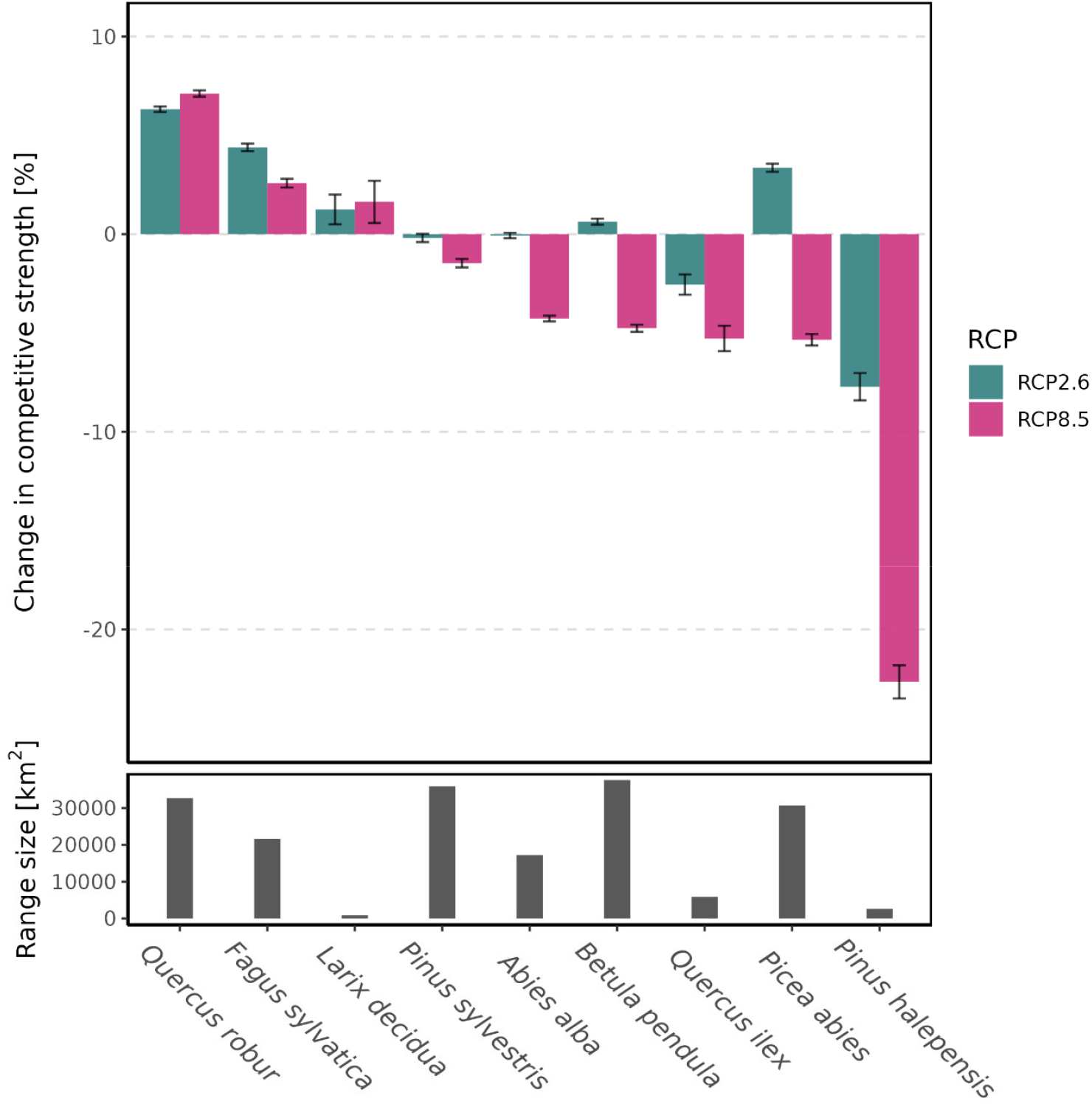
Change in competitive strength between current climate (1981 – 2010) and future climate conditions (2071 – 2100) in different climate scenarios. Values represent the mean change in competitive strength index (CSI) across all grid cells (12 x 12 km) within a species’ current distribution (area indicated in the lower panel), with error bars indicating 95% confidence intervals. CSI aggregates across the competition indicators LAI (i.e., the ability to form dense canopies and shade out other species) and height growth (i.e., the ability to overgrow competitors) by means of averaging. Detailed results for the individual components of CSI, as well as values for scenario RCP4.5 can be found in the supplementary information (Fig. S1, S2 & S3).

### Spatial patterns of changing competitive strength

Spatial variation was high among and within species, but competitive strength generally decreased in warmer and more water-limited biomes (e.g., the Mediterranean and temperate broadleaved biomes) and increased in more cold-limited biomes (e.g., temperate coniferous and boreal forests as well as tundra) (Fig. 2). European beech, which had an overall increasing competitive strength throughout its distribution, decreased in competitive strength in the Mediterranean parts of its range (−4.4%) while strongly increasing in the temperate coniferous biome (+17.5%) (scenario RCP8.5; Fig. S4). Norway spruce, showed an overall decreasing CSI throughout its distribution, which was particularly pronounced in the temperate broadleaved biome (−16.7%). Conversely, it increased in competitive strength in cold-limited areas such as Fennoscandia (+15.8% in boreal forests and +13.8% in the tundra biome). Competitive strength decreased in 79% of the current range of Aleppo pine (the species with the most pronounced negative response throughout its range), but only in 28% of pedunculate oak (the species with the most pronounced positive response), with values of 38% and 52% for European beech and Norway spruce, respectively. To determine the geographical variation in competitive responses within species, we investigated changes in CSI in different parts of a species’ climatic niche. Overall, species responded with increasing competitive strength close to their cold-induced niche edges, while close to the warm niche edges competitive strength generally decreased (Table 1). Coniferous species responded more negative than broadleaved species at their warm niche edges, and were particularly sensitive at their warm and dry niche edges.

**Table 1.** Climate-induced change in competitive strength at the edges of the current climatic niche of a species compared to the niche core. Negative values are shown in bold. Niche edges were defined as areas where mean annual temperature and annual precipitation sum were above the 90^th^ and below the 10^th^ percentile of the values of full niche space of a species, respectively. The niche core was the remaining area of a species’ niche. Values are averages of CSI and standard errors of the mean CSI, calculated from the pooled standard deviation for each group. Values for RCP4.5 are show in the Supplementary Information (Table S3).

| Niche position | RCP2.6 |  | RCP8.5 |  |
| --- | --- | --- | --- | --- |
|  | Broadleaved | Coniferous | Broadleaved | Coniferous |
| Core | 2.1 ±0.0 | **-1.1 ±0.1** | **-0.3 ±0.1** | **-7.2 ±0.1** |
| Cold-dry edge | 13.2 ±0.3 | 11.0 ±0.2 | 11.6 ±0.4 | 5.0 ±0.3 |
| Cold-wet edge | 6.0 ±0.3 | 3.3 ±0.3 | 6.0 ±0.3 | 3.0 ±0.4 |
| Warm-dry edge | **-0.3 ±0.2** | **-3.3 ±0.3** | **-2.1 ±0.2** | **-7.8 ±0.3** |
| Warm-wet edge | **-3.2 ±0.9** | **-3.2 ±0.8** | **-5.7 ±0.9** | **-4.7 ±0.9** |

**Figure 2.**
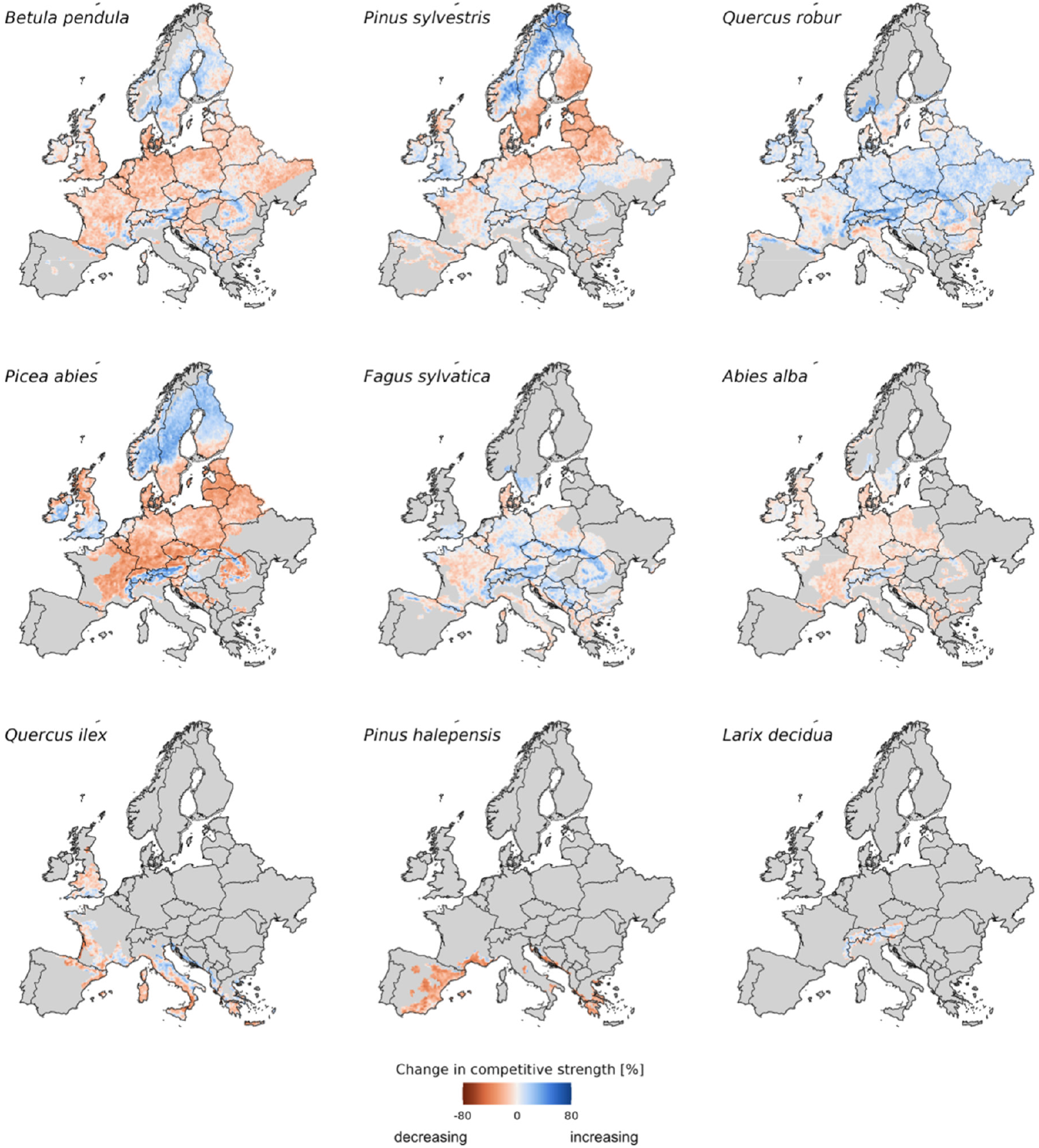
Change in competitiveness under severe climate change (RCP8.5) for nine major European tree species across their current distribution. Maps are in order of decreasing range size from top to bottom. Colors indicate the range of competitive strength index CSI changes from strongly decreasing (red) to strongly increasing (blue) CSI for the period 2071-2100 relative to 1980-2010. Maps for other climate scenarios and for the two individual components of CSI are shown in the supplementary information (Fig. S5 – S12). Uncertainty maps for CSI predictions are also available in the supplementary information (Fig. S13 – S16).

### Shifts in dominant species

A change in competitive strength only results in tree species change if another species can gain in relative competitiveness and outperform the previously dominating species. Here, we used our DNN meta-model to investigate where shifts in the currently dominant species (defined as a species having >66% of stand basal area) are likely under climate change, we found that Norway spruce, holm oak and silver birch were the species that most frequently lost their current role as dominant species to another species. Under the RCP8.5 scenario, these species were projected to lose dominance in 15.1% (7,489,178 ha), 3.9% (444,753 ha), and 3.8% (483,863 ha) of the area they currently dominate (Fig. 3a). In contrast, pedunculate oak and European beech were overall able to increase their dominance in +8.8% (4,967,027 ha) and

**Figure 3.**
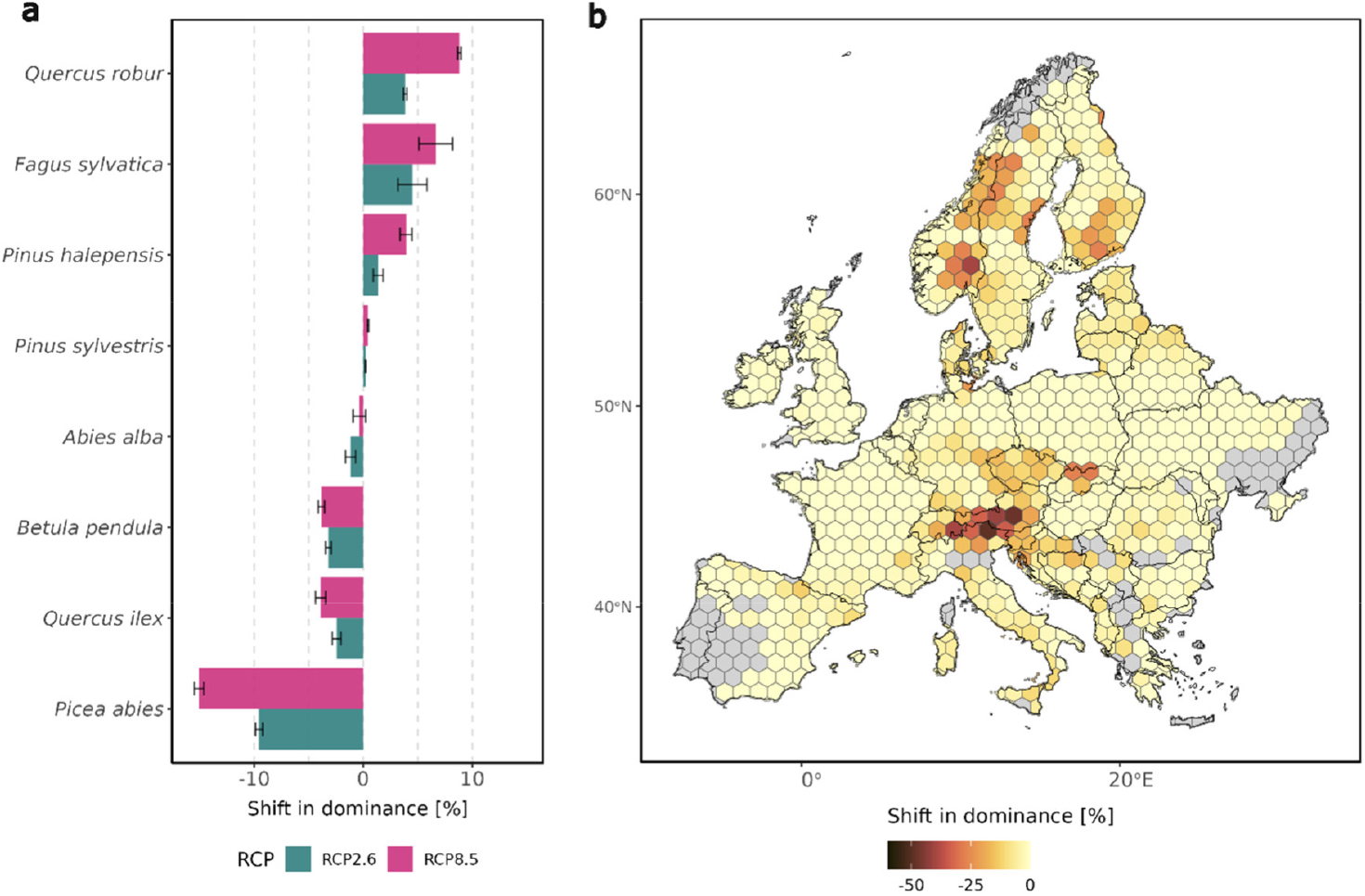
a) Losses and gains of the ability to dominate stand development under climate change across the area where a species currently dominates ^25^. b) Hotspots of change in the currently dominant species under climate change (severe climate change scenario RCP8.5, see Fig. S17 for RCP4.5 and Fig. S18 for RCP2.6). Results were aggregated from 12 x 12km grid cells to hexagons with 100 km short diagonal length (8,660.25 km^2^) for display purposes. Grey areas show regions where either no predictions were available or where none of the nine species analyzed here currently dominate. European larch was excluded from this analysis because of missing initial states (see methods for details).

+6.7% (1,386,889 ha) of the area, respectively (Fig. 3a). These projected changes in species losing and gaining dominance are largely congruent with the changes in individual-species competitive strength reported above, suggesting that changes in the competitive strength of species are also likely to result in a change in species assemblages. An exception was Aleppo pine, which was projected to moderately increase in dominance under both RCP2.6 (+1.4%; 167,655 ha) and RCP8.5 (+3.9%; 481,056 ha), despite decreasing competitive strength. This suggests that in the very warm and dry region where it occurs, its competitors may suffer even more strongly from climate change, allowing Aleppo pine to sustain its presence and even increase in dominance ^24^.

We found that on 19% of Europe’s forest area where one of the study species is currently dominant, the dominant species is likely to change under the severe climate change of RCP8.5 due to a changed competitive balance between species. This corresponds to a forest area of 30 million hectares across Europe potentially affected by changes in species dominance (15% or 24 million hectares under the moderate climate change of RCP2.6). Hotspots of expected competition-induced changes in the dominant tree species were largely located at major ecotones between biomes. One hotspot was in central Europe at the ecotone between the temperate broadleaved and temperate coniferous biomes (e.g., in the mountain regions of central Europe, Fig. 3b). Additional hotspots were at the ecotone between the tundra and boreal forests biomes (e.g., along the Scandes), as well as the ecotone between the boreal and the temperate coniferous biomes (in southern Fennoscandia). Also at the ecotone between the Mediterranean and temperate broadleaved biomes (e.g., in southern France and the Adriatic coast) elevated likelihoods for a shift in the dominant species were recorded.

## Discussion

Harnessing a novel AI-based approach, we here present the first continental-scale synthesis of local forest simulations under climate change. Mining more than 135 million simulation-years of data for more than 13,000 unique locations throughout Europe, we show that the competitive strength of major European tree species will change considerably under climate change. Our analyses highlight that evergreen coniferous tree species, currently dominating 56% of the forest area of Europe, will lose competitiveness under climate change. Conversely, deciduous broadleaved species can partly increase their competitiveness, suggesting a major shift in the dominant clades of tree species in Europe. This finding is generally in line with previous works based on statistical species distribution models ^18,26,27^. However, our process model-informed deep neural network projects rates of tree species change that are lower than those inferred from correlation-based species distribution models. While species distribution models predict equilibrium states under given climate conditions and do not consider competition between species explicitly, the process-based models underlying our analyses simulate the interactions between trees as emergent property based on local resource availability and tree species traits ^19,20^. The change in dominant species inferred from our synthesis of process-based simulations is thus likely a more realistic indicator of species change and underlines that major shifts in tree species are particularly expected at the trailing edge of the distribution of individual species. This is in line with already observed trends at the warm range edges of important European tree species ^28–30^. We note that our assessment of where species dominance could change uses current species distribution as reference, and economically important tree species such as Norway spruce were historically cultivated outside of areas where they would dominate naturally.

Our AI-based approach offers a novel solution to bridging the gap between detailed local-scale forest modeling and continental-scale analyses of forest dynamics under climate change. Traditional process-based models, while excellent for capturing fine-scale processes such as the competitive interactions between trees (e.g., ^31,32^), face computational limitations when applied across large spatial scales. Furthermore, these models are usually developed and parameterized for specific focal ecosystems, trading off broad-scale applicability across a diverse range of ecosystems for locally accurate projections. Our AI-driven synthesis leverages the strengths of these local models while overcoming their limitations by effectively scaling their responses to the continental scale. Beyond the intricacies of each individual model, our AI-based approach distills the emergent responses to environmental drivers, thus resulting in a more robust projection of forest dynamics overcoming the individual shortcomings of models. Furthermore, our approach allows the assimilation of future local model simulations to further refine simulated continental-scale responses, facilitating collaborative research on forest ecosystems in a changing world.

We here explicitly considered height growth and LAI as indicators of competitive strength. Both indicators are specifically relevant in the context of competition for light, with height growth being a crucial strategy in the asymmetric competition of plants for light ^33^ and high LAI indicating the ability to shade out competitors ^34^. While our choice of indicators reflects the fact that light is the dominant constraint for tree growth in large parts of Europe ^35^, we note that other traits such as rooting depth and the tolerance of extremes (e.g., drought, frost) are also important factors for the competitiveness of trees ^36,37^. We found that LAI generally responded more negatively to climate change than height growth and that only European beech was able to increase LAI while all other species were not. This finding contrasts large-scale simulation results with earth system models and dynamic vegetation models, generally expecting a greening of Europe under climate change ^38,39^. As recent observations in Europe are not in line with a general greening of vegetation ^40^, the findings presented here underscore the value of our novel, AI-based process-model synthesis. Future work could harness the approach pioneered here to further elucidate where and under which conditions large-scale earth system models differ from best-available local simulation approaches, helping to quantify uncertainties and improve global modeling capabilities ^41,42^.

Inherent limitations of our approach need to be considered when interpreting our findings. Some parts of Europe – both in geographic and climatic space – remain underrepresented in the process model simulations considered here. Expanding the simulation database with simulations from these regions (e.g. parts of Eastern Europe, but see Fig. S13 – S16), could enhance the accuracy and reduce uncertainties in our predictions. Furthermore, while an important strength of our approach is the synthesis across many different process models – hedging against uncertainties in the structure and process representation of individual models – the data underlying the current study are not equally distributed across models and species ^23^. For instance, training data for Aleppo pine was obtained from a single process-based model, rendering the strong climate change response reported here more uncertain compared to other, more broadly represented species. We also note that extreme climatic events and natural disturbances such as wildfire, wind and bark beetle outbreaks are important drivers of tree species change ^43,44^; yet, these factors were not considered in our analysis. Addressing such processes explicitly should thus be a future direction of research. Furthermore, changes in dominant tree species reflect changes in the relative competitiveness between species. We studied nine species in detail, and included a total of 63 species in our training dataset for the DNN model of dominance. Nonetheless, including an even broader set of species in process model simulations would be desirable, as it could identify local winners and losers under climate change more comprehensively. We note, however, that particularly species that have historically been rare remain difficult to parameterize in process-based models ^45^.

Changes in the competitive strength of tree species and related shifts in dominant vegetation have important implications. The diversity of species communities in forests, for instance, is strongly linked to the dominant tree species ^46^. Changes in dominant tree species could thus trigger a turnover in species communities, particularly since climate change could trigger a shift from evergreen coniferous trees to deciduous broadleaved trees, i.e. species groups that diverge strongly in both morphology and phylogeny. The finding that deciduous oaks might increase in competitiveness could have a positive effect on biodiversity, as these species have been found to be associated with highly diverse species communities in previous analyses ^18,47^. Changes in the competitive strength of tree species also have important implications for managing forests for ecosystem services. In plantations for timber production, the loss of competitive strength of a crop species may necessitate increased efforts in tending and thinning. This is particularly relevant as evergreen conifer species represent the majority of plantation forests in Europe. In forests managed for multiple ecosystem functions and services, maintaining diversity in structures and species is an important management approach ^48,49^. This requires an intimate understanding of the competitive relations between trees and their potential changes under climate change. Our spatially explicit analysis of hotspots of potential future changes in dominant tree species can alert managers to areas where changes are likely in the future, highlighting situations in which adjustments of the prevailing silvicultural concepts might be needed to maintain desired tree species mixtures. We conclude that climate change will strongly alter the competitive relationships between trees across Europe. This change can have important implications for biodiversity and the supply of ecosystem services, and should thus be considered more explicitly in forest policy and management.

## Materials and Methods

### Methods overview

To investigate the impact of climate change on the competitive strength of tree species and assess related potential shifts in species dominance, we trained a Deep Neural Network (DNN) on a large dataset of harmonized forest simulations across Europe ^23^, containing data on 135 million simulation-years for 13,599 unique locations throughout Europe. We subsequently applied the trained DNN to predict forest state changes under different climate scenarios, analyzing a set of indicators of competitiveness at the level of individual species, as well as the DNN-projected changes in species composition (Fig. 4). In a first step, harnessing the dataset of harmonized forest simulations, we converted continuous forest simulation outputs to discrete forest states (based on tree height, LAI, and species composition, see ^50^ and coupled them with data on climate and environmental conditions to create training data for deep learning. In a second step, we trained a DNN to predict transitions between these states based on driver data (i.e., current state of the system, climate scenario data). Informed by local simulations from the underlying process-based models, the trained DNN served as meta-model to project climate-induced changes (cf. ^21^. We used this meta-model to predict changes in the three state variables for nine study species under current and future climate conditions across the current distribution range of the species. Based on these results we investigated how competitive strength (represented by the indicators height growth and LAI) and dominant tree species shifted under future conditions. Fig. 4 gives an overview of the workflow of the analysis, all steps are explained in more detail below.

**Figure 4.**
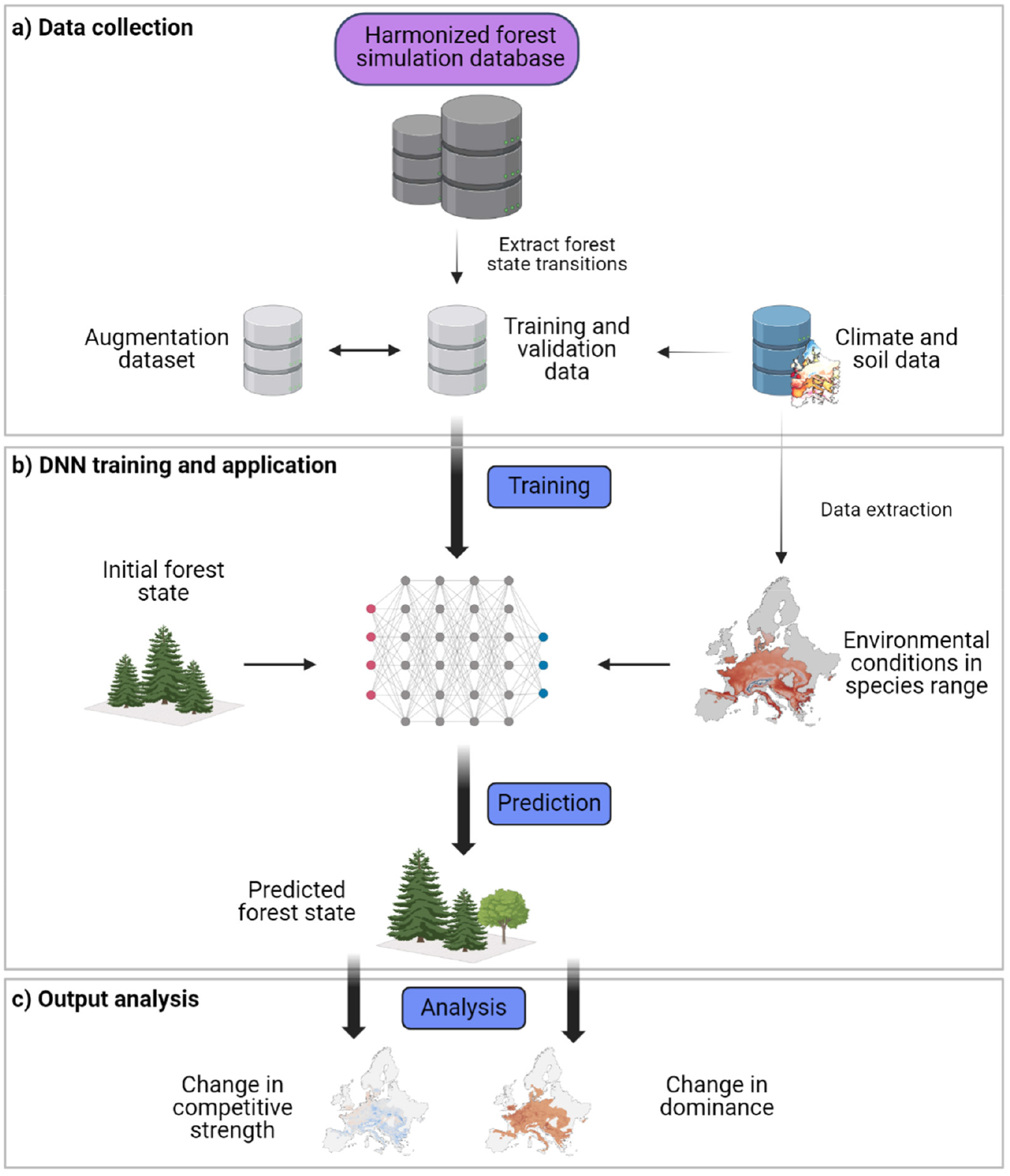
Analysis workflow. a) Data collection from a harmonized database of process-based model simulations under climate change ^23^. From the database, forest state transitions were extracted to create training and validation datasets, using a data augmentation approach to increase the number of training data. In parallel, corresponding data on environmental conditions (i.e. climate and soil) were compiled from pan-European datasets to enable linking forest state transitions to environmental conditions. b) DNN training and application: training data was used to train a DNN to predict (i) the forest state after a transition and (ii) the time until a transition takes place, based on the initial forest state and the prevailing environmental conditions. c) Output analysis: we analysed differences between predictions under baseline climate (1981-2010) and future climate (2071-2100) for indicators of competitive strength (objectives i) and ii)) and changes in the dominant tree species (objectives iii) and iv)). This figure was created with BioRender.com.

### Data on forest dynamics

We here used for the first time a recently published harmonized database of more than 135 million simulation-years from 1.1 million simulation runs conducted by 17 different models in 13,599 locations across Europe ^23^. This database is a collection of previously conducted local forest simulations under climate change, which were harmonized to a common suite of output variables and standardized metadata (including a vector of common environmental drivers). All simulations in the dataset are based on process-based models operating at the stand-to landscape-scale. Models were run in the absence of natural disturbance, and either represent unmanaged conditions or business-as-usual management for a given location and species. Management signals were filtered out during the training dataset preparation (see below). The various output structures of the different forest models were harmonized to three common variables: Canopy height, leaf area index, and species composition. These three variables were chosen as they describe complementary aspects of forest ecosystems and as they were available from all underlying models.

We converted continuous model outputs to discrete states by binning the three output variables, in order to simplify the classification task for the DNN ^50^. Classes for species composition described the dominant and up to four co-occurring species. A species was considered *dominant* when the proportion of basal area was higher than 66% of the total basal area, other species were considered as admixed species when they exceeded 20% of the simulated basal area. LAI (one-sided or projected) was stratified into three discrete classes: 0-2 m^2^/m^2^, 2-4 m^2^/m^2^, and >4 m^2^/m^2^. Canopy height was binned in 2m classes, ranging from 0-2m to 48-50m, and >50m. The unique combination of classes in all three dimensions resulted in a potential maximum of 4.1 million forest states. However, only a small fraction of these potential states is actually realized in ecosystems, as not all combinations occur in reality.

In a next step, we identified transitions between these states from the forest simulations in the database. A transition was detected whenever the forest state changed from one year to the next in the underlying simulations. Additionally, we identified ‘no-change’ conditions, i.e. situations where the same state persisted for at least 10 years. State transitions were the primary response variable used for training the DNN; consequently, we henceforth refer to them as ‘training samples’. Training samples also contained auxiliary information on the environmental conditions under which a state transition occurred. The final training sample comprised information on the forest state prior to transition, its residence time (i.e., the time elapsed since the last transition), the forest state after transition, and the time when the next transition is expected to happen, alongside soil conditions and climate data for the time horizon of the transition. In other words, each training sample contained the current state of a forest as well as information about whether, how, and when the state will change within the next decade, given the prevailing soil and climate drivers.

We expanded our dataset by creating augmentation training samples based on the original training samples. Data augmentation is a widely used and well-established technique to enhance the number of training data for DNNs ^51^, enabling models to generalize better without overfitting. We created augmentation training samples leveraging the state transitions from the original training samples. Specifically, we altered the residence time and time until transition by the same number of years, maintaining the absolute duration between state transitions. By doing so, we preserved the integrity of the temporal dynamics of the underlying data, ensuring that the total time for each state transition remained unchanged. This method allowed us to increase the number of training samples meaningfully, without artificially generating new state transitions or introducing redundancy.

We filtered training samples by location, model and forest state to reduce the bias resulting from multiple contributions from the same model and location to the simulation database. Specifically, the majority of simulations in the database are from two models (i.e. iLand and 4C), and some species were simulated more frequently than others ^23^. To reduce the dominance of a single model and/ or species in the training dataset we downsampled to a maximum of 100,000 training samples per location, while keeping as many different forest states as possible. Furthermore, we filtered out direct management signals from interventions such as thinnings or final harvests (e.g. by detecting and eliminating canopy height reduction by more than 2 m from the data) from the training samples, yet indirect management signals might still be included in the data. The resulting dataset comprised 2,750,456 million data points covering 5,445 distinct forest states. The number of training samples for the different species ranged from 34,759 (*Betula pendula*) to 250,634 (*Fagus sylvatica*; see Table S1).

### Climate and soil data

Pan-European climate change scenarios and soil information were used (i) to provide environmental context information for the state transitions from the underlying simulation database ^23^, and (ii) as driver data for model projections. As environmental context information for state transitions we used daily climate data for mean temperature, precipitation, solar radiation and vapor pressure deficit (VPD). We aggregated daily data to annual averages for all four variables and also derived monthly averages for temperature and precipitation. Furthermore, we used a machine learning-based targeted compression to derive a set of climate indices relevant for predicting climate responses of vegetation. This step considerably reduced the amount of data required in processing (by a factor of 30), but retained the information value of high temporal resolution climate data for climate impact simulations. A detailed description of the climate data compression is given in the supplementary methods (section *Climate compressor approach*). The final climate data for each training sample contained mean annual temperature, mean annual precipitation, mean annual solar radiation, mean annual VPD, monthly means for temperature and precipitation, and 24 climate indicators from the climate data compression. (i.e. in total 52 variables) for ten years. As driver data for model projections, we used climate data for two time slices, baseline climate (1981-2010) and future climate (2071-2100) in 10-year timesteps. We used EURO-CORDEX daily climate data for three Representative Concentration Pathway (RCP) scenarios (RCP2.6, RCP4.5, RCP8.5) as well as historical climate conditions, each simulated with three Global Circulation Models (GCMs) (for details see supplementary methods section *Climate data*). We bias-corrected climate data based on ERA-Interim (ECMWF) data. All climate data was obtained in a 0.11° x 0.11° (∼12 x 12km) spatial resolution from the Copernicus Climate Data Store. (https://cds.climate.copernicus.eu/cdsapp#!/dataset/projections-cordex-domains-single-levels?tab=overview).

We compiled pan-European soil datasets at a 1 x 1km resolution to represent the variability of soil conditions within each climate grid cell. For soil depth, soil texture (i.e. sand, silt and clay content) and water holding capacity we obtained gridded data from the European Soil Data Center (Hiederer, 2013). Furthermore, we approximated plant-available nitrogen based on a pseudo-mineralization rate derived from a continental-scale spatial model in combination with a continental product of soil N stocks from the SoilGrids dataset ^53^. For more details see ^23^.

### Deep neural network architecture and evaluation

DNNs are powerful tools to learn complex relationships in data and are increasingly used in the environmental sciences ^54,55^. The DNN architecture used here consisted of a feed-forward neural network with 6.6 million trainable parameters, arranged in 22 layers with 3 blocks with residual connections. The inputs of the DNN were: the present forest state, the history of the forest state (i.e. the three previous forest states of a cell), its residence time, the residence time history (i.e. the residence times in the previous three states), soil conditions, and climate conditions. The DNN was trained to classify the forest state after a transition and the time until transition, considering 5,445 discrete forest states and ten classes of target time (10-year forecasting window). We used TensorFlow ^56^ in combination with the Keras API ^57^ to implement the model architecture. To check the ability of the DNN to generalize we performed several cross-validation experiments, including a random five-fold cross-validation, a model selection cross-validation and a climate scenario cross-validation. As expected, data are distributed unequally between states, with many states occurring only rarely while some occuring very frequently. We thus further tested the predictive performance of the DNN relative to state frequency to ensure that our model does not predominantly predict the most frequent states (for details see supplementary methods section *DNN architecture and cross-validations*). Moreover, we analyzed the importance of individual variables in projections using a variable permutation technique (Figure S20). The final DNN performed well, correctly predicting the forest state after transition with 86.9% and the time to transition with 61.1% accuracy in the validation dataset (Table S2).

### Prediction of forest state transitions with the trained DNN

We used the trained DNN for predicting forest state transitions for the nine study species. To assess competitive strength (objectives i) and ii)) we focused on the stem exclusion stage of stand development (i.e., the stage in which competition is highest) as we assumed that changes in competitiveness will be most relevant in this stage of stand development. Initial states for prediction were defined as canopy heights in the stem exclusion stage (i.e. between 15-30% of maximum tree height; obtained from ^18^), and simulations were started from all three LAI classes. To account for the local variability in soil conditions (1 x 1 km), three soil vectors were sampled for each 12 x 12km grid cell. To analyse shifts in tree species dominance (objectives iii) and iv)), we used the 10 most common forest states in the dataset in which the focal species currently dominates stand development (>66% of basal area) as starting point, again focusing on the stem exclusion stage of stand development. We excluded European larch from this part of the analysis because no forest states with admixed species were available for the stem exclusion stage in our simulation database. Only forest states with a minimum of 10 occurrences in our dataset were considered as initial states, ensuring sufficient representation for prediction.

Projections for all species were done for all grid cells within their current range. Species ranges were obtained from the chorological maps of European woody species ^58^. We combined native and naturalized ranges whenever species were occurring outside of their native range and the information was available. We obtained model predictions for each species with several initial states, for six 10-year time steps (three baseline, three future time steps), three GCMs and three RCPs. For each species, model predictions were averaged across the three different timesteps of a period (baseline or future), different initial states and soil conditions. For each of the three forest state variables (canopy height, LAI, and species composition) differences were calculated between simulations under climate change and those under current climate, with the latter serving as counterfactual to isolate climate change effects.

### Competitive strength

Height growth and LAI were used as indicators for competitive strength. We calculated height growth (i.e. change in canopy height) and change in LAI from DNN-derived state transitions. Specifically, we divided the simulated state change by the predicted time until transition. To overcome the uncertainty of time to transition predictions in the analysis, we used a weighted mean (with prediction probabilities as weights) of the three transition times that were predicted as most probable by the DNN. Changes in the two indicators per grid cell were calculated as differences of percent negative and percent positive changes under climate change relative to projections under historic climate. To aggregate across both dimensions we calculated a competitive strength index (CSI) per tree species as the mean change in height growth and LAI across all grid cells comprising the range of a species. To investigate a species’ response to climate change at its range edges, we distinguished the core range (between the 10^th^ and 90^th^ percentile of mean annual temperature and mean annual precipitation of the current range of a species) from the niche edges (beyond the 10^th^ and 90^th^ percentile, respectively) in temperature and precipitation space, considering the warm-dry, warm-wet, cold-dry and cold-wet edge of the distribution. CSI was again calculated as average across all grid cells within the respective part of the niche.

### Species dominance

To assess whether changes in competitive strength actually result in a change in the dominant tree species, we analyzed the species composition component of our DNN projections. Specifically, we analyzed whether and how frequently a species that currently dominates stand development (i.e., >66% of basal area) loses its dominant position under climate change. We expressed this shift in dominance in percent change under climate change relative to baseline climate conditions. To map hotspots of change in the dominant tree species across Europe, we extracted all grid cells in which one of our nine study species currently dominates stand development (based on ^25^). Note that for Aleppo pine and holm oak we masked their distribution range with the area where *miscellaneous pine*, respectively *miscellaneous oaks* were mapped as dominant, due to the lack of species level information. We subsequently analyzed the percentage of predictions per grid cell in which the dominant species was likely to lose its dominance under climate change, aggregating to hexagons with 100 km short diagonal length (area of 8,660.25 km^2^) for a better visual identification of hotspots of tree species change. To avoid overemphasizing individual negative predictions, we set a minimum threshold of at least 5% prediction per grid cell with a negative shift in dominance in order to report dominance loss.

### Uncertainty estimates

All DNN predictions were probabilistic, i.e. probabilities for the predicted future forest state were obtained from the DNN, and the most probable state analyzed further in the context of our research questions (see above). To further elucidate uncertainties of our DNN meta-model, we examined the probability of the predicted forest state transitions and times to transition across the range of each species (Fig. S13 – S16). These maps indicate the degree of certainty that the DNN has in a projected state transition for a given cell. They highlight that for species where high amounts of training data were available (i.e. European beech, Norway spruce, Scots pine) the DNN tended to be more confident than for species with less training data. The spatial variation in the thus obtained uncertainty maps suggests that regions poorly covered by underlying simulation data had higher uncertainty in DNN predictions. For instance, predictions for silver birch in Eastern Europe had lower confidence due to the limited availability of training data for that species in this region.

## Data and code availability

Data will be published on a Zenodo repository upon acceptance. For reviewers, the data is available here: https://syncandshare.lrz.de/getlink/fiNYL4ugJMrog6vNivYtE8/. Code for the data preprocessing, deep neural network, network predictions and the analysis is available at https://github.com/magrueni/europe_tree_competition_publi

## Acknowledgments

We acknowledge funding received from the European Union’s Horizon 2020 research and innovation program under grant agreement no. 101000574 (RESONATE: Resilient forest value chains – enhancing resilience through natural and socio-economic responses). R.S. and W.R. acknowledge further support from the European Research Council under the European Union’s Horizon 2020 research and innovation program (Grant Agreement 101001905, FORWARD).

