## Supplementary Material for "Loss of competitive strength in European conifer species under climate change"

### Supplementary information

This file contains supplementary material for the manuscript entitled *Loss of competitive strength in European conifer species under climate change*. It contains supplementary material (Tables S1 & S3; Figures S1-S18) and supplementary methods (including Figures S19-S22).

**Supplementary tables:**

Table S1: Study species and number of datapoints in the training and validation dataset. Maximum tree height calculated from Wessely et al., 2024. We obtained the optimal growing phase heights as 15-30% of the maximum tree height and rounded the values to match to the 2m canopy height classes.

| Species | Number of datapoints in dataset | Maximum tree height [m] | Optimal growing phase [m] (rounded to the next 2m height class) |
|---|---|---|---|
| *Fagus sylvatica* | 250,634 | 45 | 6-14 |
| *Pinus sylvestris* | 248,319 | 45 | 6-14 |
| *Picea abies* | 247,298 | 60 | 10-18 |
| *Quercus robur* | 191,269 | 45 | 6-14 |
| *Abies alba* | 119,212 | 65 | 10-20 |
| *Larix decidua* | 102,130 | 50 | 8-16 |
| *Pinus halepensis* | 79,718 | 25 | 4-8 |
| *Quercus ilex* | 62,916 | 25 | 4-8 |
| *Betula pendula* | 34,759 | 30 | 4-10 |

Table S2: Summary table of the Deep Neural Network (DNN) performance metrics. We tested accuracy of the model predicting the correct label and predicting the true label in the Top 2 predictions. The shown numbers are the mean and standard deviation (SD) of the 5-fold cross-validation (CV) with five randomly sampled equal size chunks of training data. The DNN was trained five times, each time using one chunk as validation set while the other four were used for training. Two more cross-validation experiments are shown in the supplementary information.

| | **Training set** | | **Validation set** | |
|---|---|---|---|---|
| Output | target state | target time | target state | target time |
| Final DNN | 0.904 | 0.672 | 0.869 | 0.611 |
| Final DNN Top 2 | 0.974 | 0.854 | 0.954 | 0.793 |
| CV Mean Correct | 0.889 | 0.658 | 0.864 | 0.608 |
| CV Mean Top 2 | 0.968 | 0.838 | 0.953 | 0.788 |
| CV SD Correct | 0.0004 | 0.0012 | 0.002 | 0.0031 |
| CV SD Top 2 | 0.0001 | 0.0008 | 0.0006 | 0.0044 |

Table S3: Climate-induced change in competitive strength at the edges of the current climatic niche of a species compared to the niche center. Negative values are shown in bold. Niche edges were defined as areas where mean annual temperature and annual precipitation sum were above the 90th and below the 10th percentile, respectively. The niche core was the remaining area of a species' niche. Values are averages of CSI and standard errors of the mean CSI, calculated from the pooled standard deviation for each group.

| | **RCP2.6** | | **RCP4.5** | | **RCP8.5** | |
|---|---|---|---|---|---|---|
| Niche position | Broadleaved | Coniferous | Broadleaved | Coniferous | Broadleaved | Coniferous |
| Core | 2.1 ±0.0 | **-1.1 ±0.1** | 0.6 ±0.1 | **-3.2 ±0.1** | **-0.3 ±0.1** | **-7.2 ±0.1** |
| Cold-dry edge | 13.2 ±0.3 | 11.0 ±0.2 | 8.3 ±0.4 | 12.8 ±0.2 | 11.6 ±0.4 | 5.0 ±0.3 |
| Cold-wet edge | 6.0 ±0.3 | 3.3 ±0.3 | 5.9 ±0.3 | 3.7 ±0.4 | 6.0 ±0.3 | 3.0 ±0.4 |
| Warm-dry edge | **-0.3 ±0.2** | **-3.3 ±0.3** | **-4.0 ±0.2** | **-7.7 ±0.3** | **-2.1 ±0.2** | **-7.8 ±0.3** |
| Warm-wet edge | **-3.2 ±0.9** | **-3.2 ±0.8** | **-4.5 ±0.8** | **-6.7 ±0.9** | **-5.7 ±0.9** | **-4.7 ±0.9** |

**Supplementary figures:**

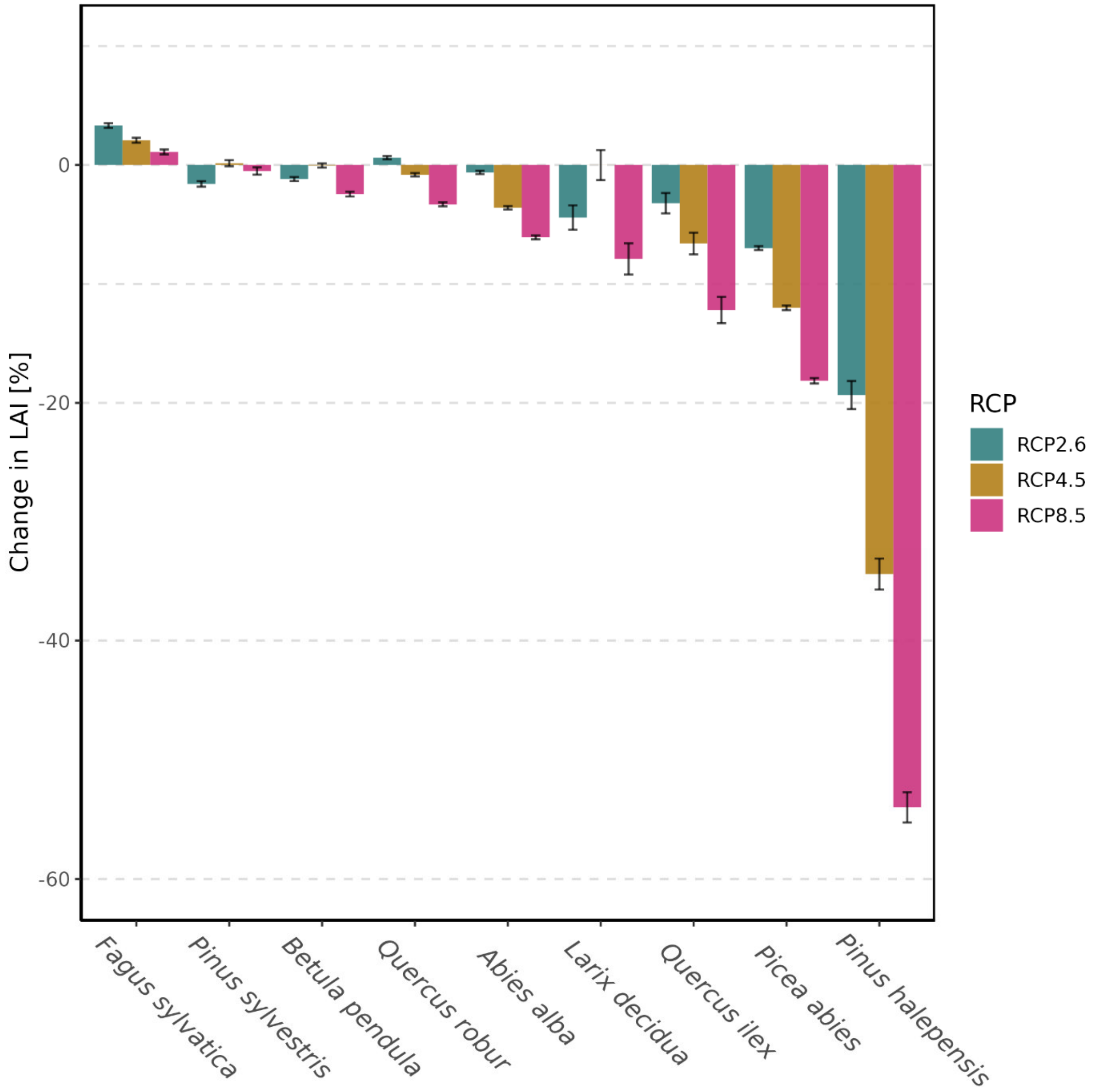


Figure S1: Change in the LAI component of the competitive strength index (CSI) between the climate of the recent past (1981 – 2010) and future climate (2071 – 2100) conditions in different climate scenarios (RCP2., RCP4.5 and RCP8.5). Values represent the mean change in LAI across all grid cells (12 x 12 km) within a species' current distribution with error bars showing the 95% confidence intervals.

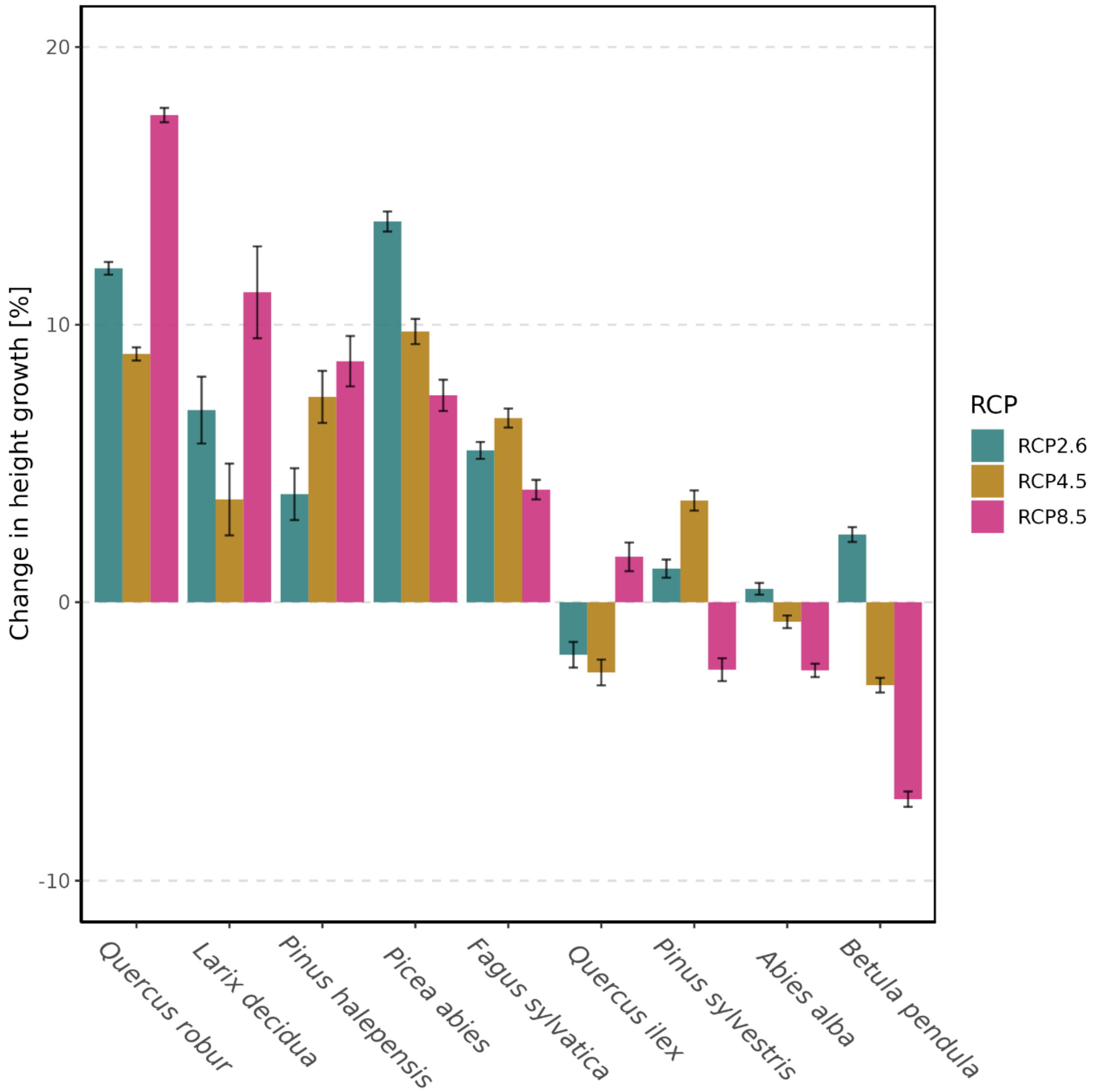


Figure S2: Change in the height growth component of the competitive strength index (CSI) between the climate of the recent past (1981 – 2010) and future climate (2071 – 2100) conditions in different climate scenarios (RCP2., RCP4.5 and RCP8.5). Values represent the mean change in height growth across all grid cells (12 x 12 km) within a species' current distribution, with error bars showing the 95% confidence intervals.

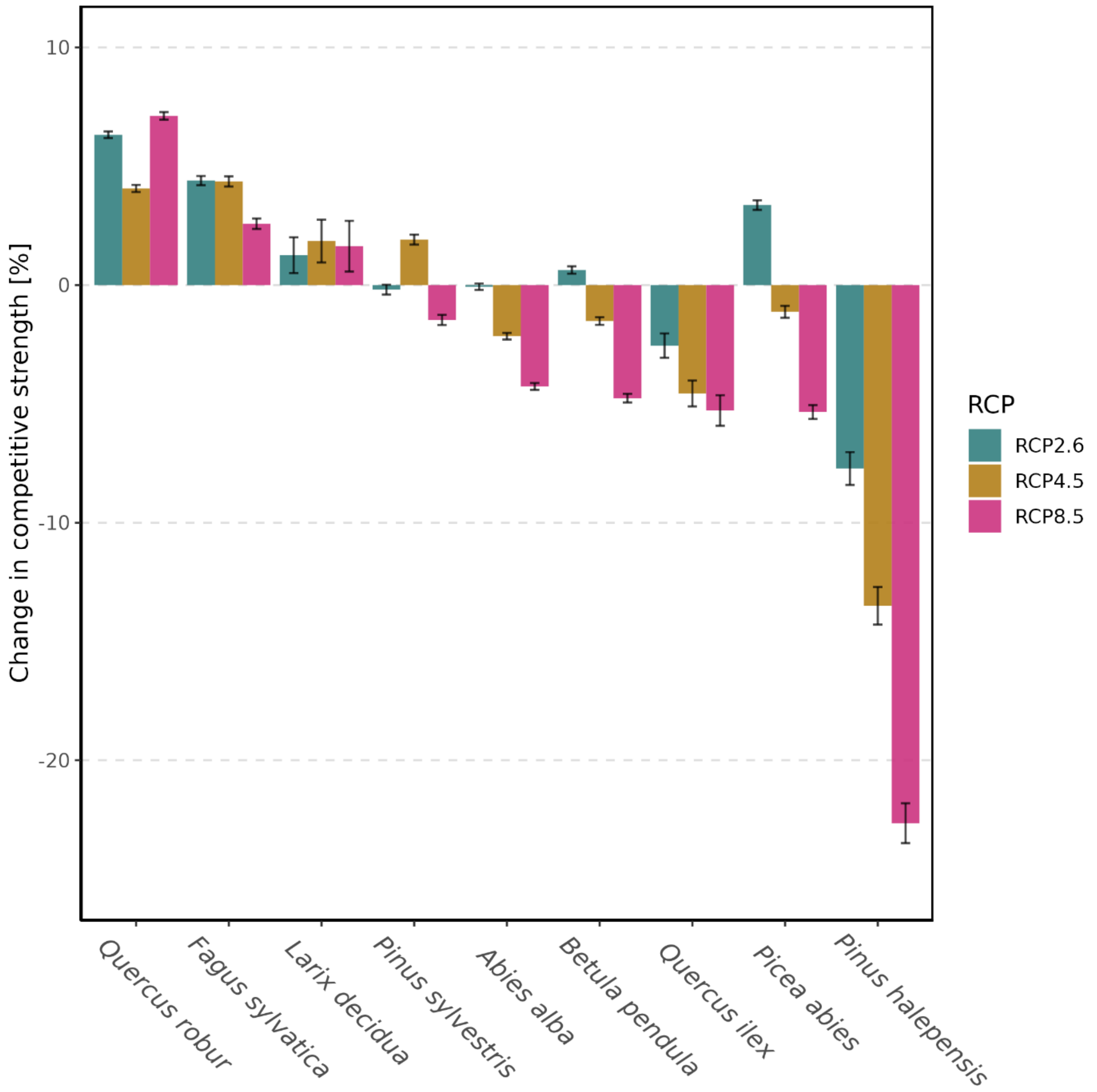


Figure S3: Change in competitive strength index (CSI) between the climate of the recent past (1981 – 2010) and future climate (2071 – 2100) conditions in different climate scenarios (RCP2., RCP4.5 and RCP8.5). Values represent the mean change in competitive strength across all grid cells (12 x 12 km) within a species' current, with error bars showing the 95% confidence intervals. CSI aggregates across the indicators LAI (i.e., the ability to form dense canopies and shade out other species) and height growth (i.e., the ability to overgrow competitors) by means of averaging.

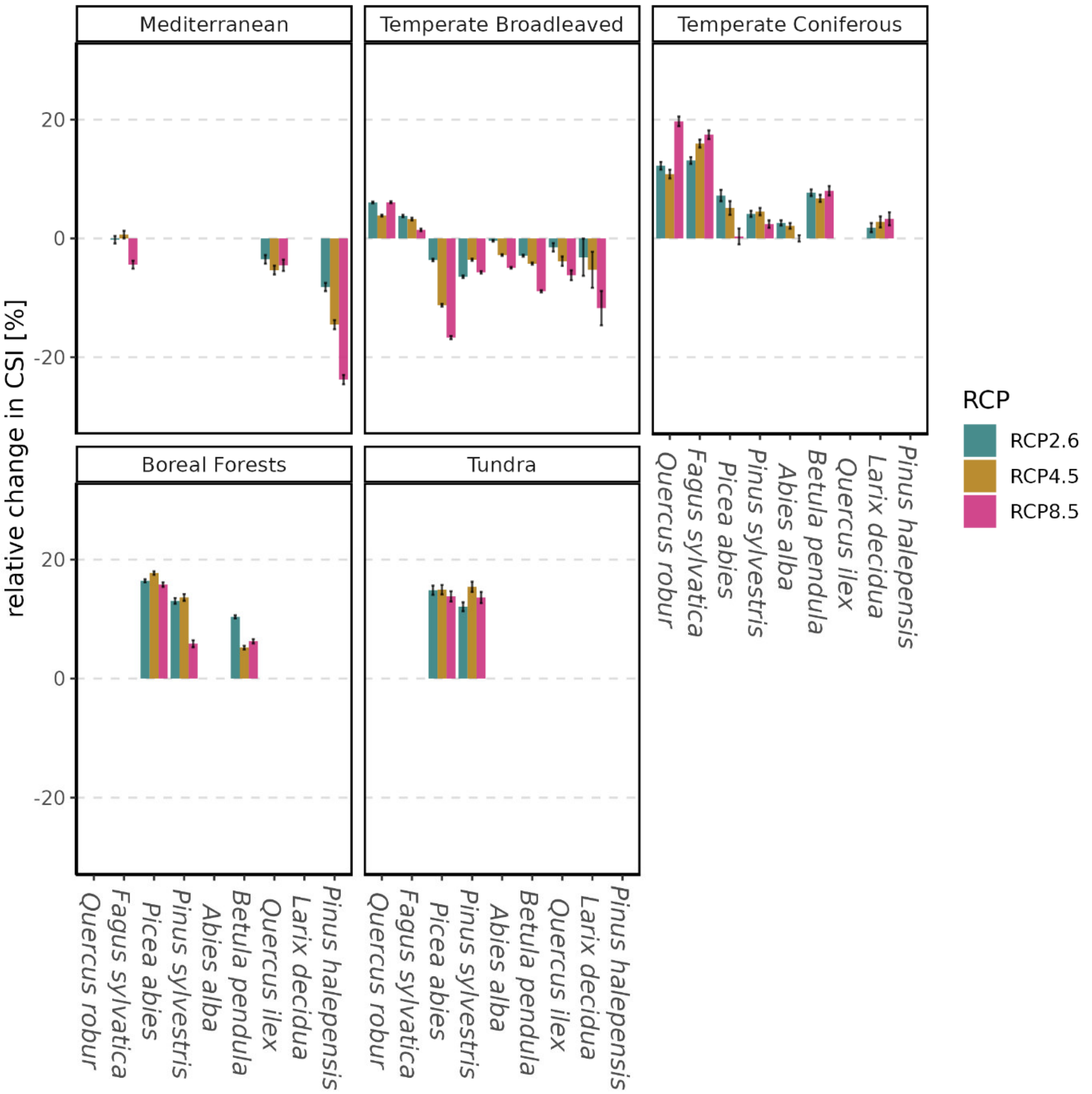


Figure S4: Change in competitive strength index (CSI) between the climate of the recent past (1981 – 2010) and future climate (2071 – 2100) conditions in different climate scenarios (RCP2., RCP4.5 and RCP8.5) in each biome separately. Values represent the mean change in competitive strength across all grid cells (12 x 12 km) within a species' current, with error bars showing the 95% confidence intervals. CSI aggregates across the indicators LAI (i.e., the ability to form dense canopies and shade out other species) and height growth (i.e., the ability to overgrow competitors) by means of averaging.

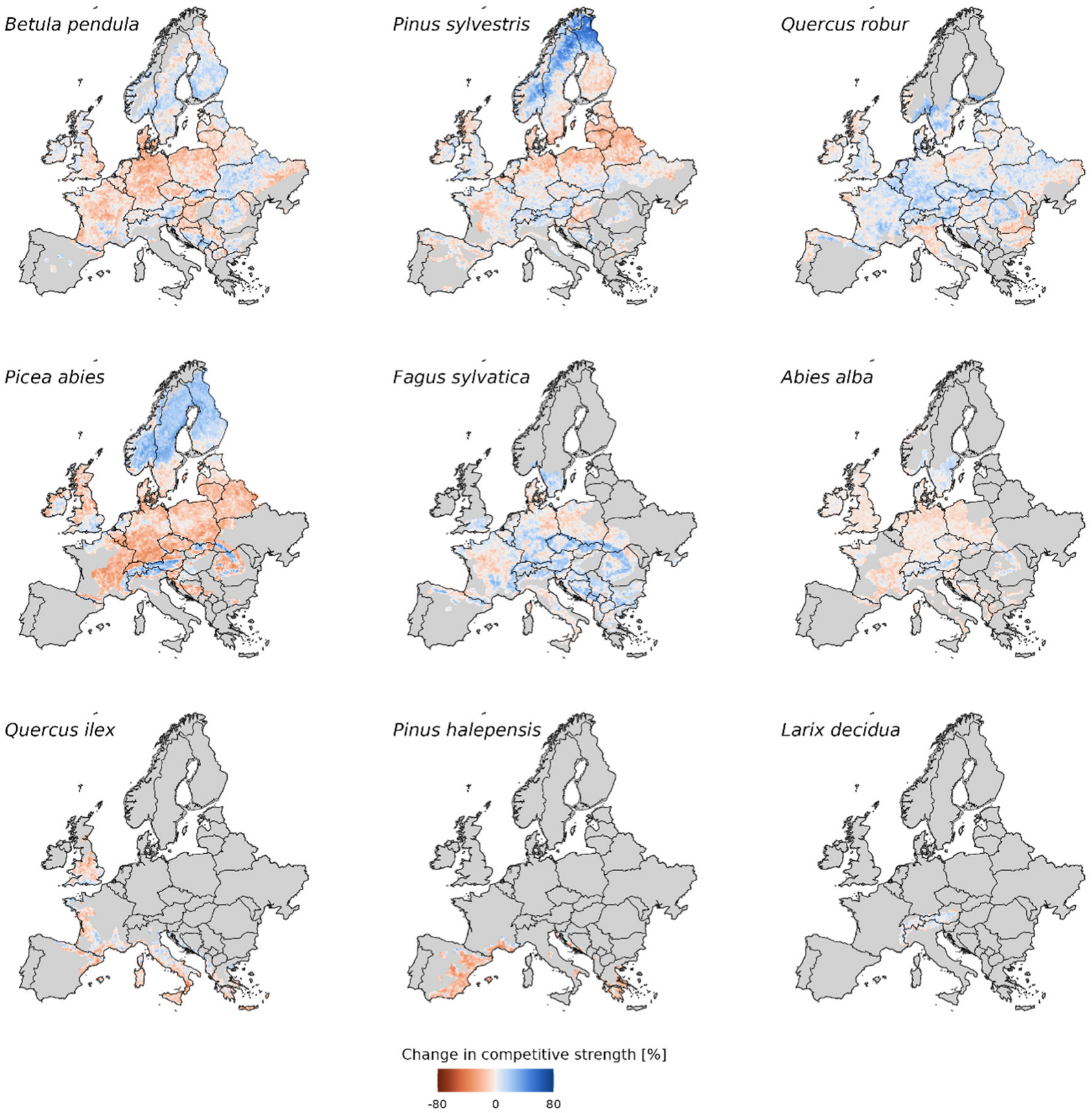


Figure S5: Change in competitiveness (competitive strength index CSI) under RCP4.5 for nine major European tree species across their distribution range. Maps are in order of decreasing range size from top to bottom. Colors indicate the range of CSI changes from strongly decreasing (red) to strongly increasing (blue) CSI.

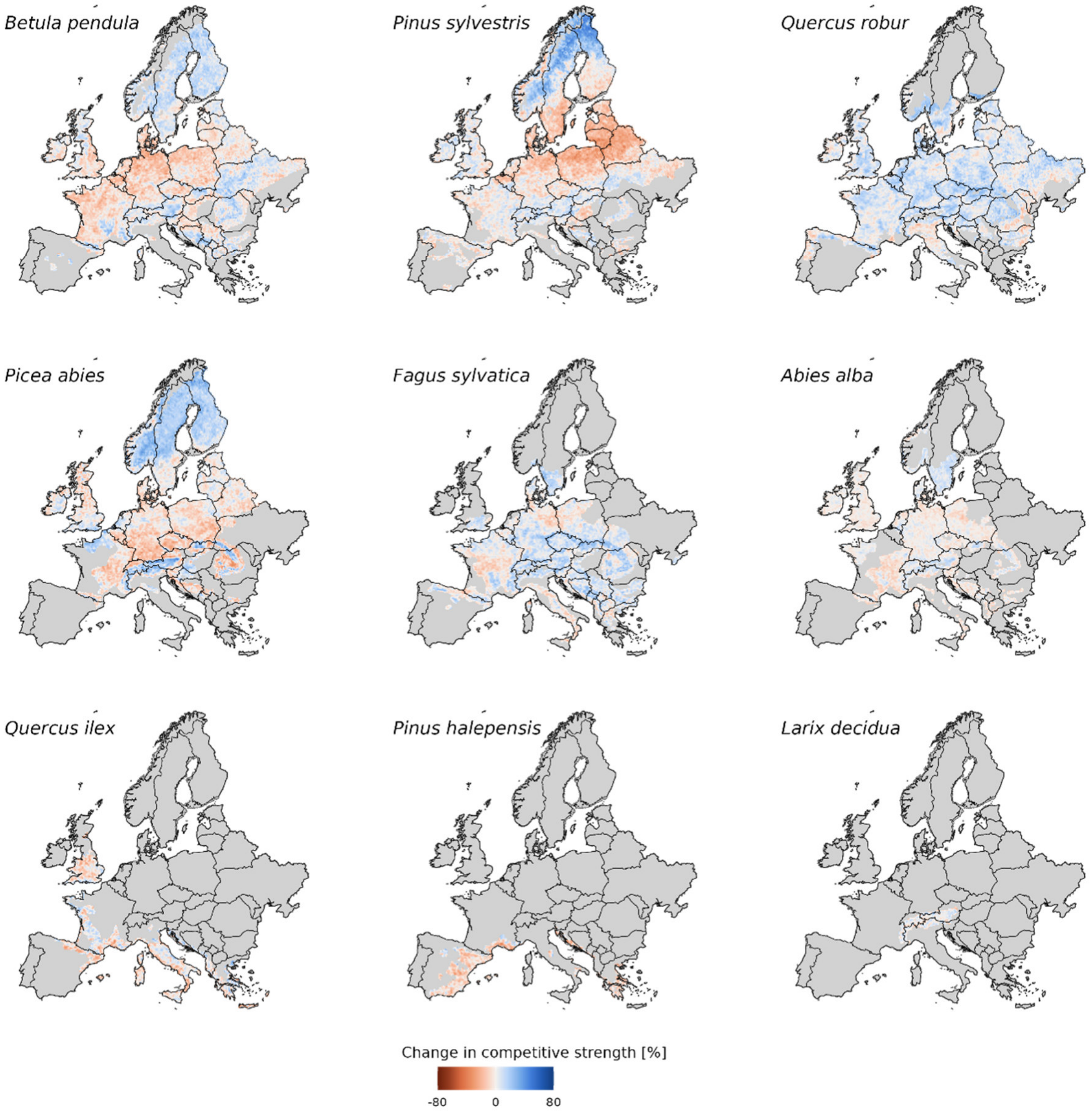


Figure S6: Change in competitiveness (competitive strength index CSI) under RCP2.6 for nine major European tree species across their distribution range. Maps are in order of decreasing range size from top to bottom. Colors indicate the range of CSI changes from strongly decreasing (red) to strongly increasing (blue) CSI.

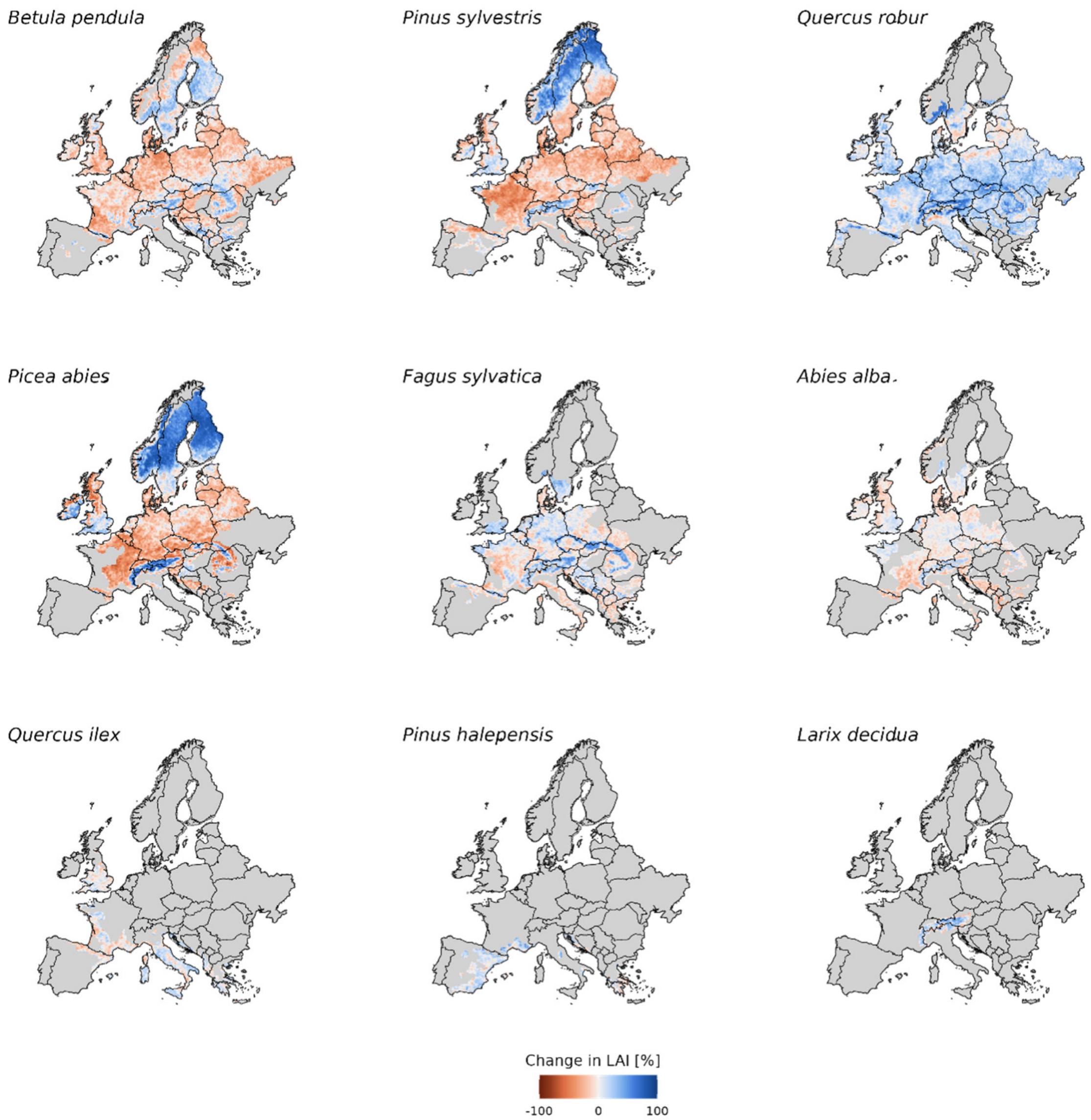


Figure S7: Change in LAI under severe climate change (RCP8.5) for nine major European tree species across their distribution range. Maps are in order of decreasing range size from top to bottom. Colors indicate the range of CSI changes from strongly decreasing (red) to strongly increasing (blue) CSI.

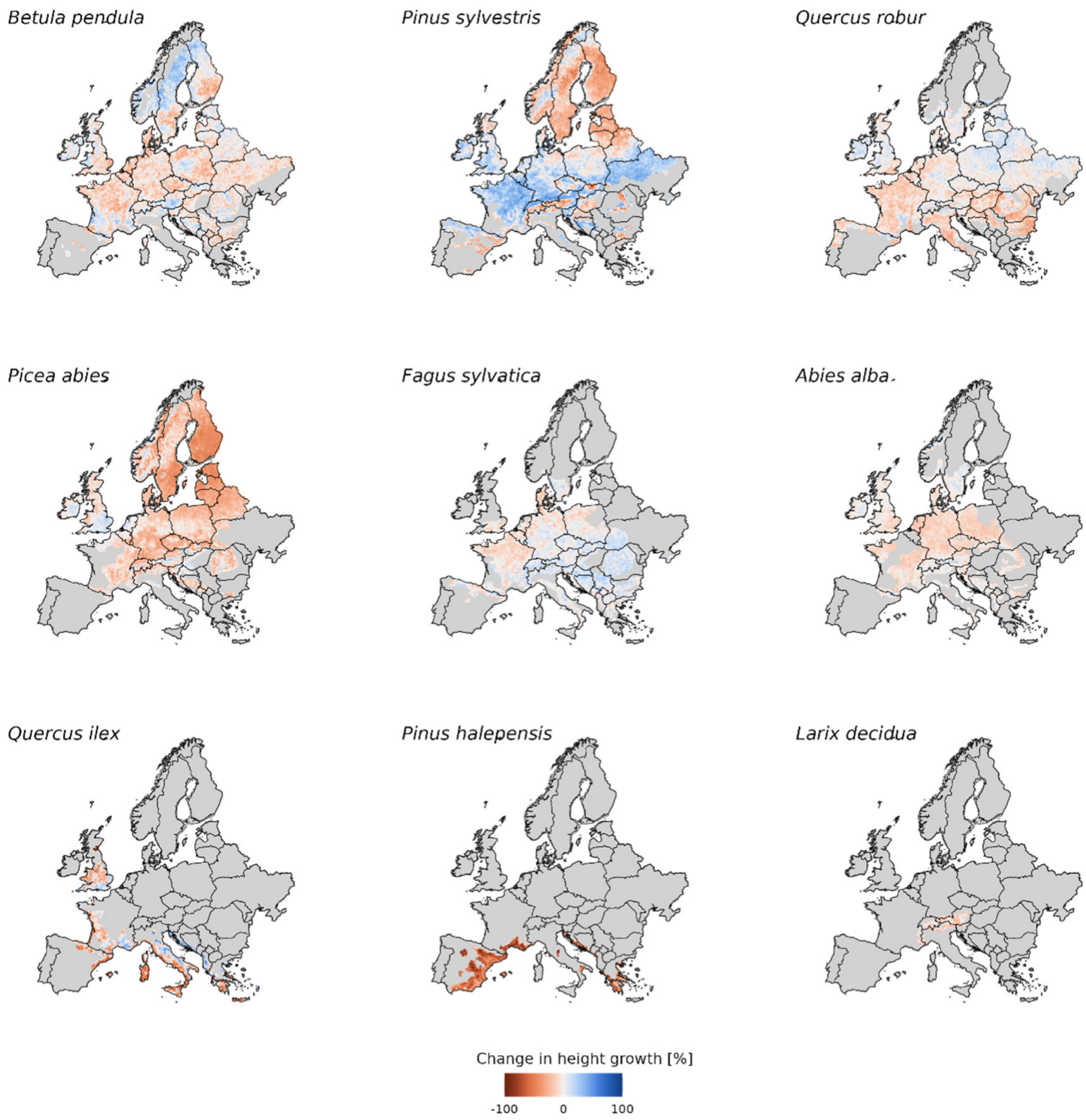


Figure S8: Same as Figure S7 but for height growth change under RCP8.5.

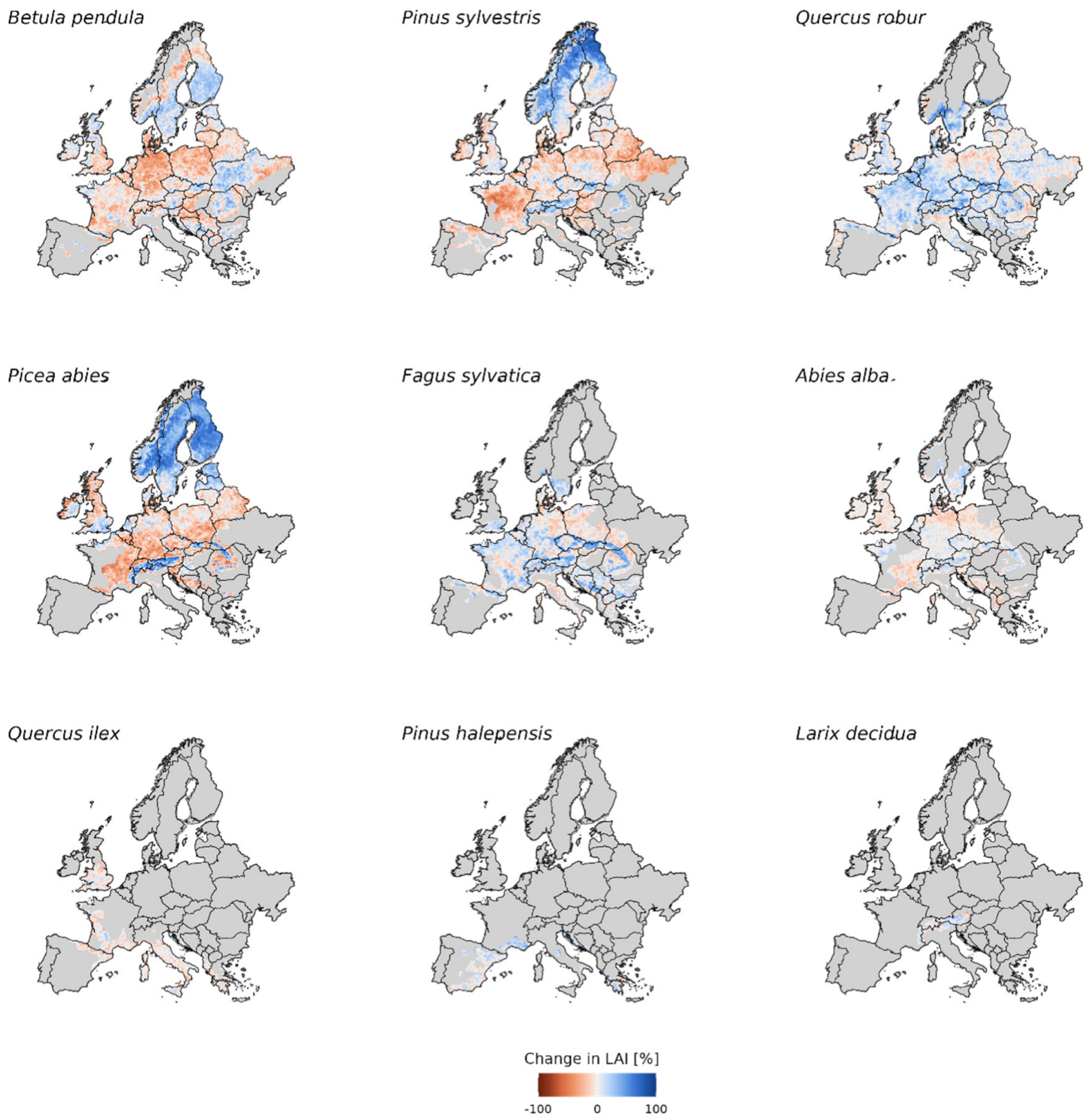


Figure S9: Same as Figure S7 but for LAI change under RCP4.5.

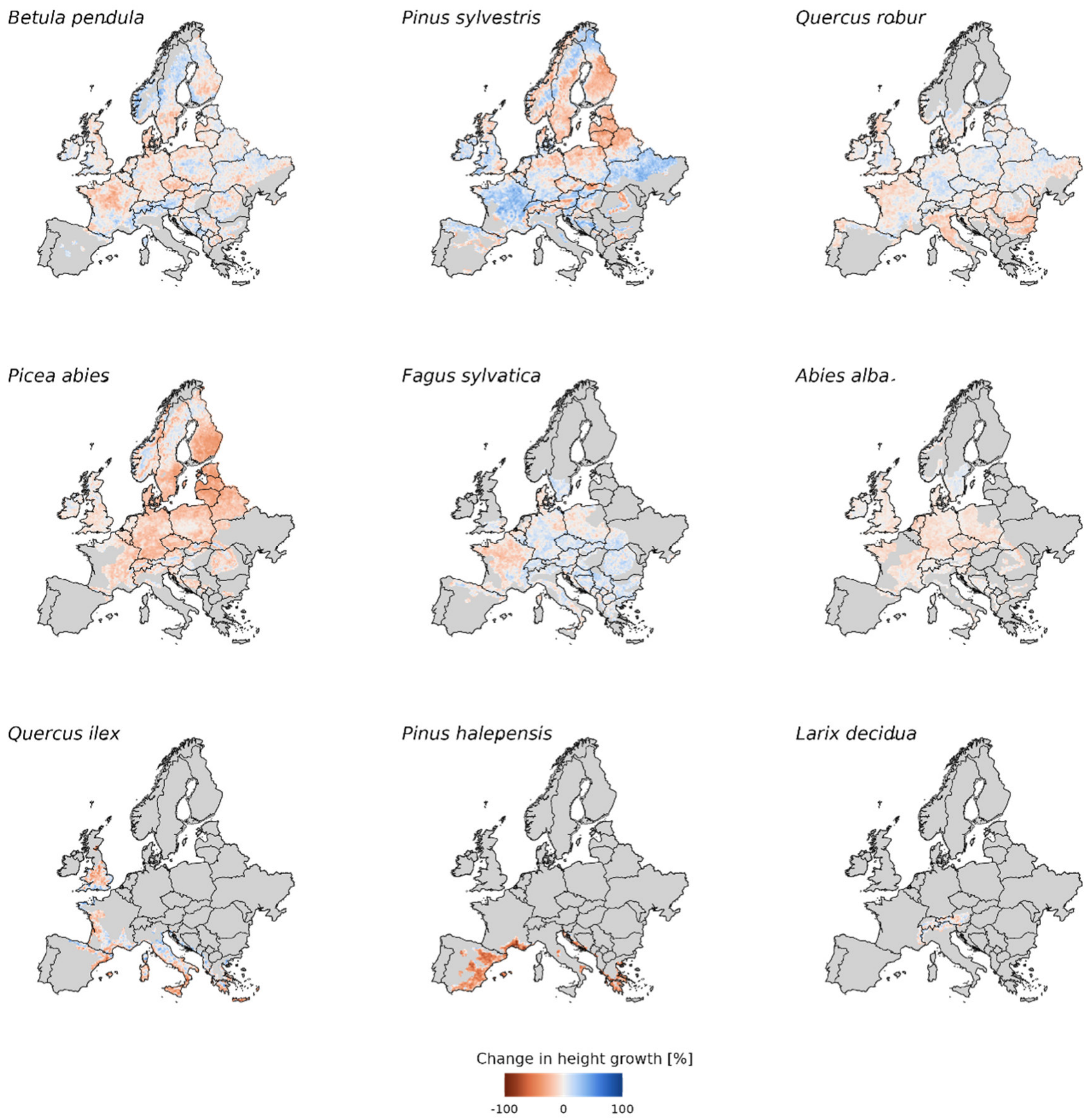


83

84 Figure S10: Same as Figure S7 but for height growth change under RCP4.5.

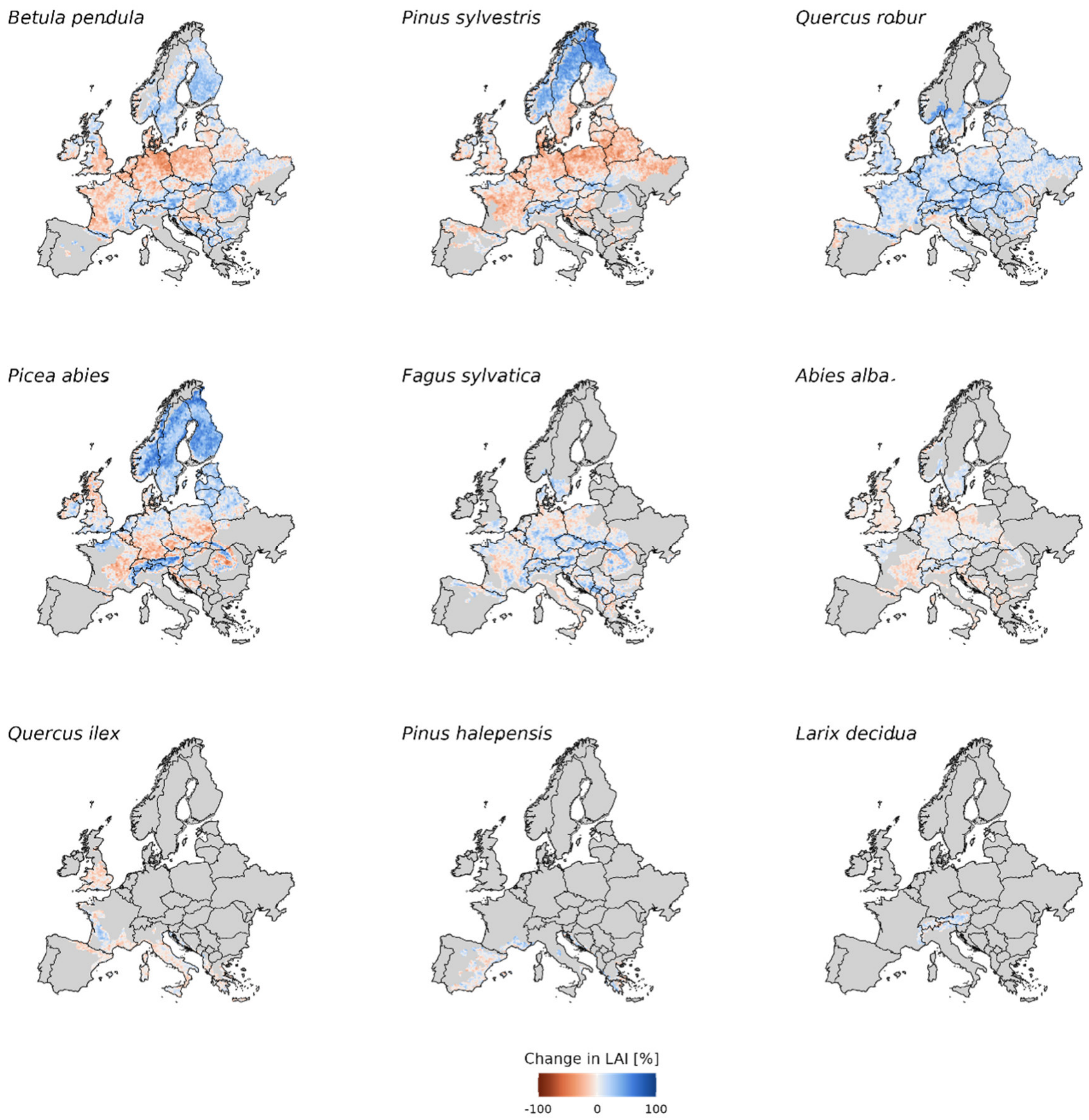


Figure S11: Same as Figure S7 but for LAI change under RCP2.6.

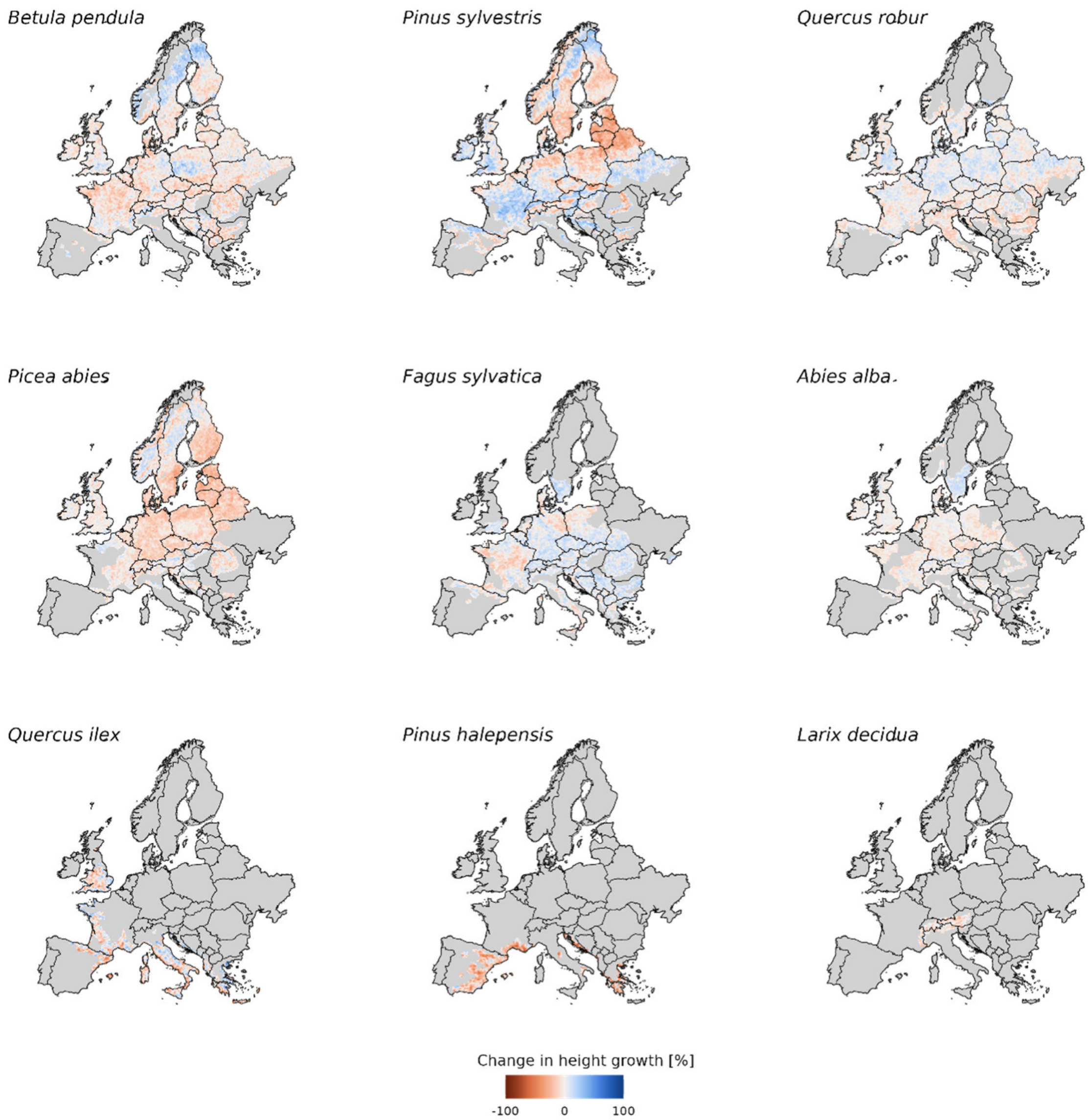


Figure S12: Same as Figure S7 but for height growth change under RCP2.6.

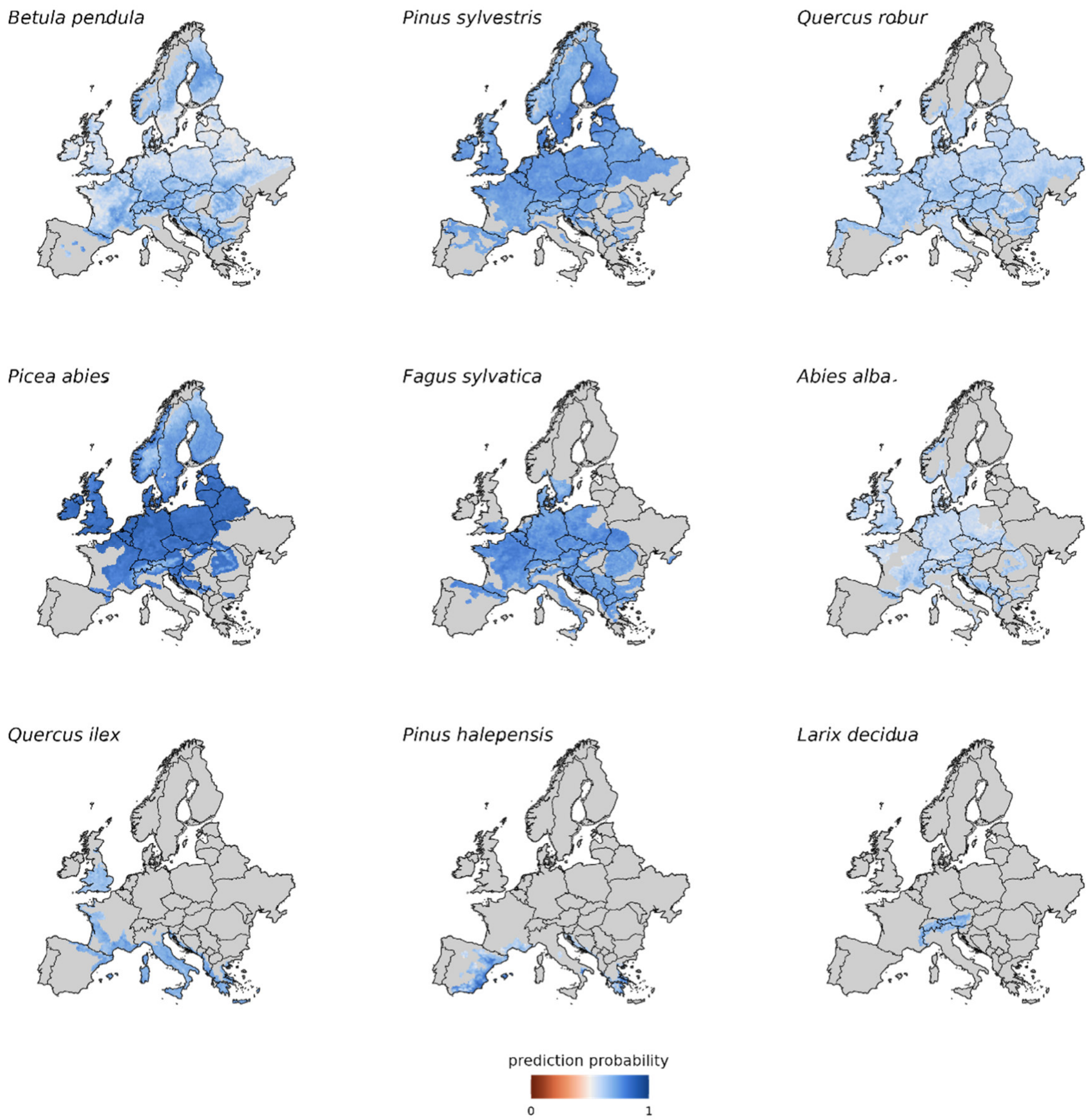


Figure S13: Uncertainty maps of spatial predictions for target states. For this we obtained the prediction probability for the predicted state by the DNN and averaged it for all predictions across GCMs and initial states per grid cell. The resulting values can be interpreted as how confident the DNN is with the predictions across geographical space for the different species. This figure shows the state prediction probabilities for predictions under baseline conditions.

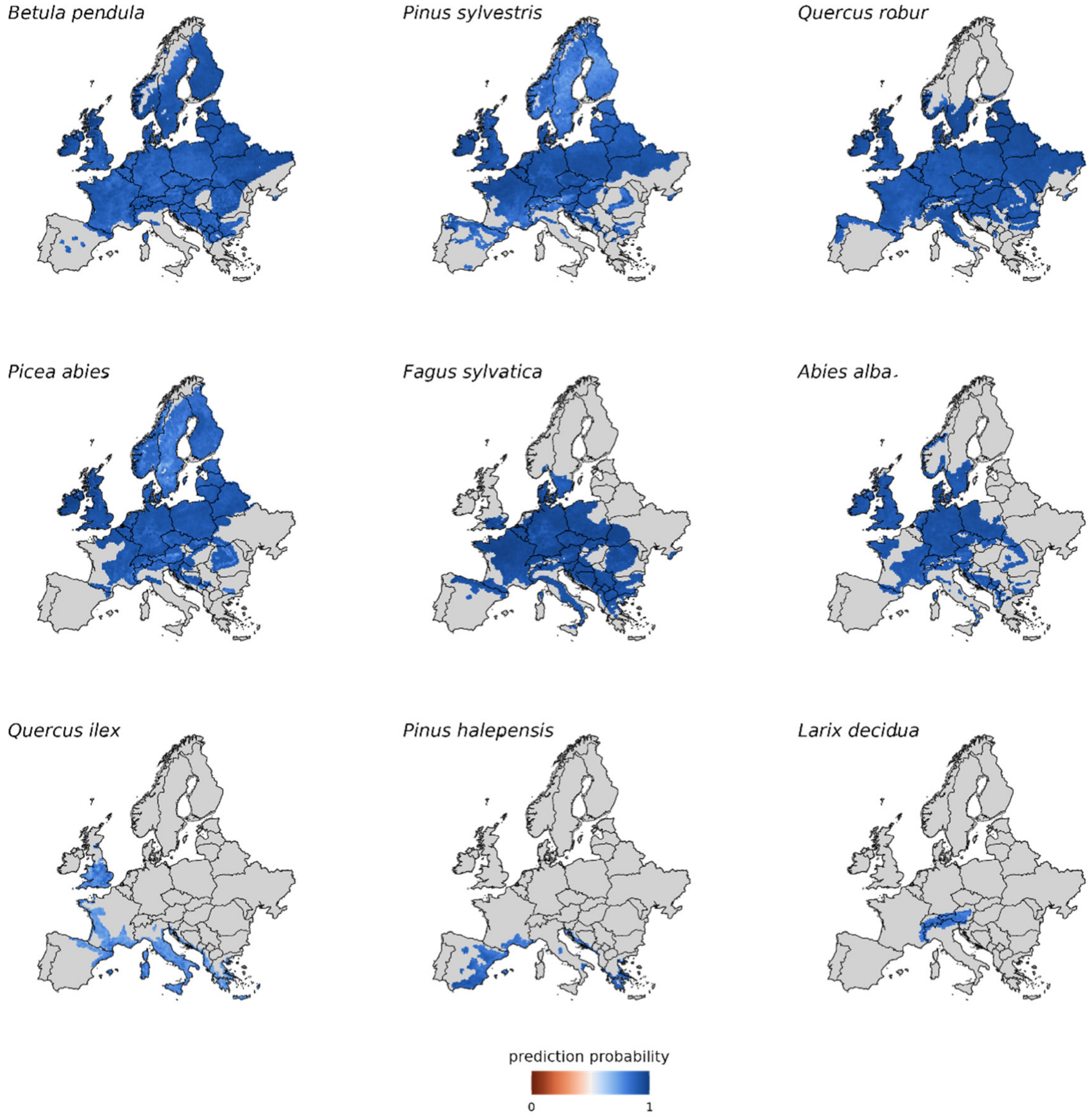


Figure S14: Same as S13 but for the predictions of target time. Note that for the analysis we used a weighted mean of the top 3 target time predictions to overcome the lower accuracy of the DNN to predict target time. Therefore, here we show the sum of the top 3 prediction probabilities.

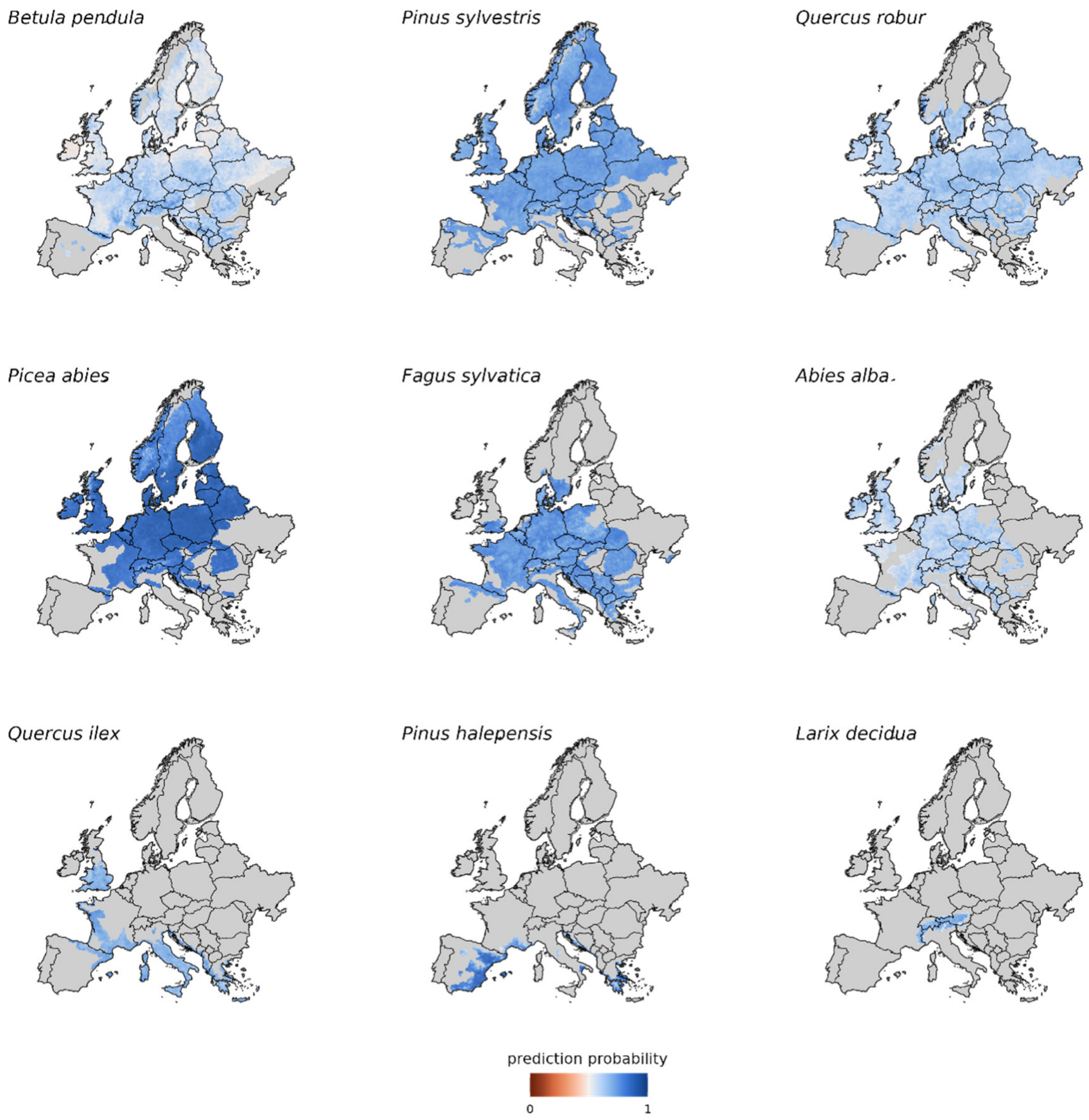


Figure S15: Same as figure S13 but for state prediction probabilities under future conditions (RCP8.5).

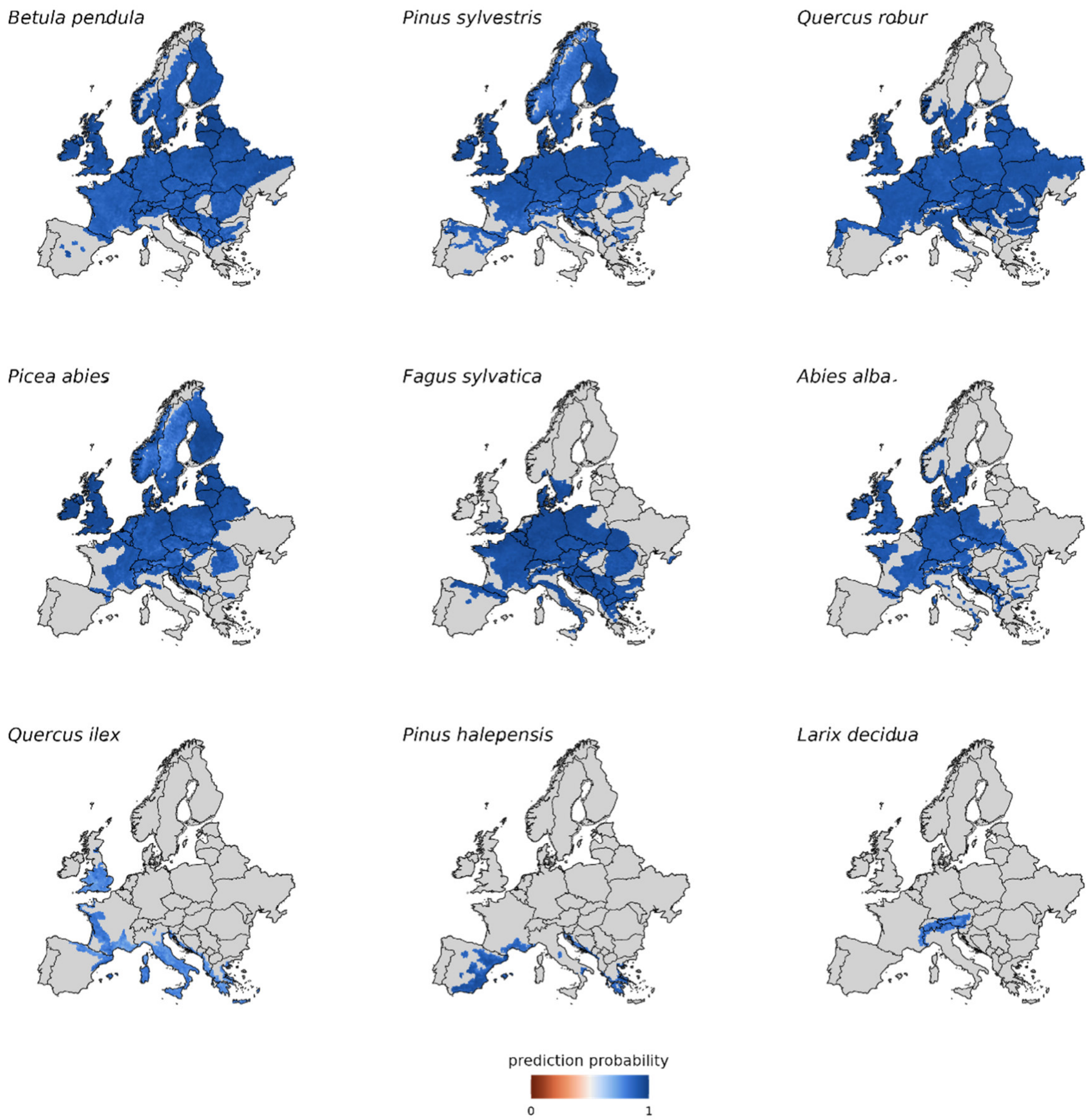


Figure S16: Same as figure S13 but for target time prediction probabilities under future conditions (RCP8.5).

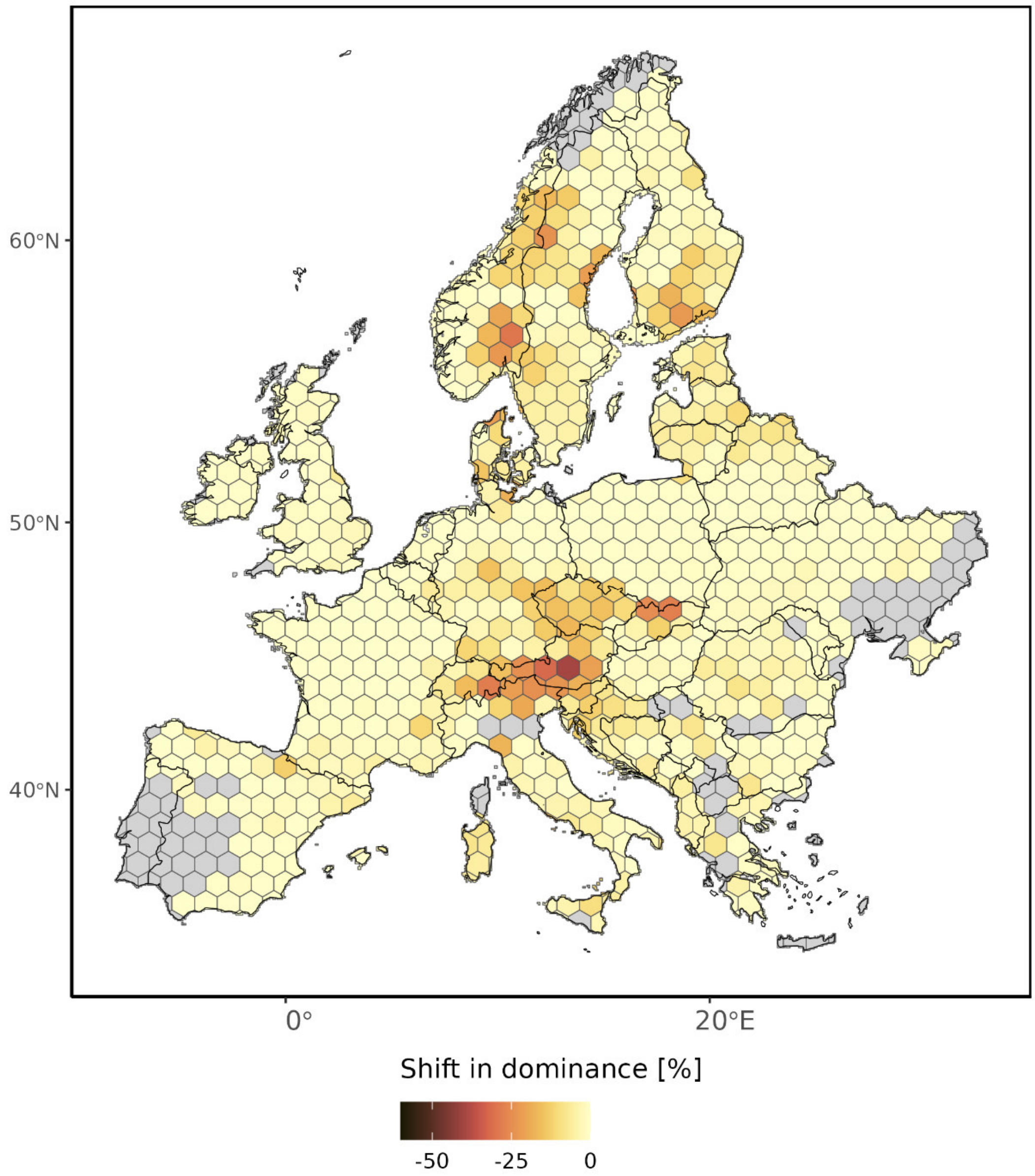


Figure S17: Hotspots of change in the currently dominant species under climate change (scenario RCP4.5). Results were aggregated from 12 x 12km grid cells to hexagons with 100 km short diagonal length (8,660.25 km$^2$) for clarity. Grey areas show regions where either no predictions were available or where none of the nine species analyzed here currently dominates. Note that European larch was excluded from this part of the analysis as there were no suitable initial states available in the database (see methods for details).

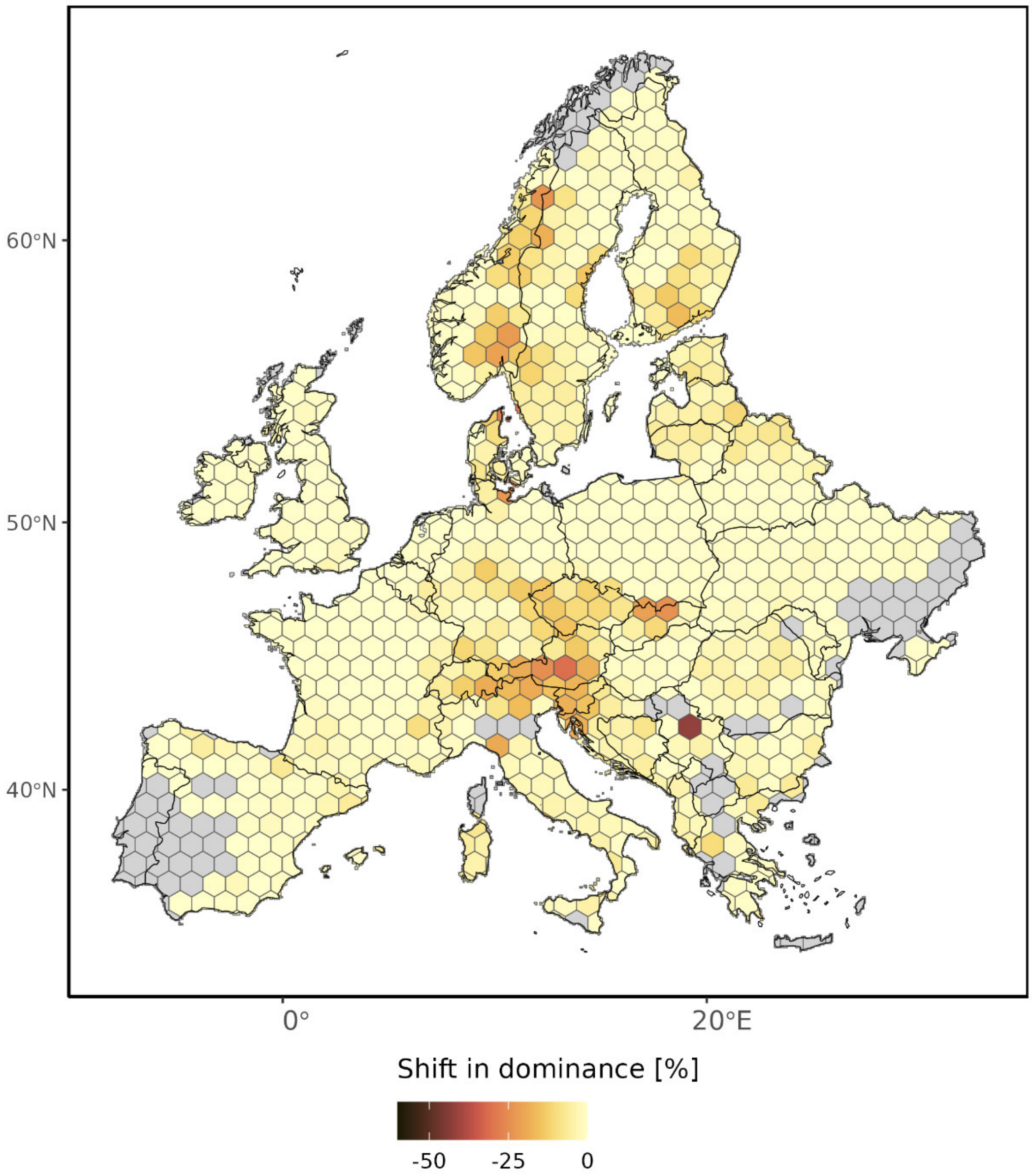


Figure S18: Hotspots of change in the currently dominant species under climate change (scenario RCP2.6). Results were aggregated from 12 x 12km grid cells to hexagons with 100 km short diagonal length (8,660.25 km$^2$) for clarity. Grey areas show regions where either no predictions were available or where none of the nine species analyzed here currently dominates. Note that European larch was excluded from this part of the analysis as there were no suitable initial states available in the database (see methods for details).

#### Supplementary methods:

*Data augmentation and filtering*

We expanded our dataset by creating augmentation training samples based on the original training samples. Data augmentation is a widely used and well-established technique to enhance the number of training data for DNNs (Shorten & Khoshgoftaar, 2019), enabling models to generalize better without overfitting. We created augmentation training samples leveraging the state transitions from the original training samples. Specifically, we altered the residence time and time until transition by the same number of years, maintaining the absolute duration between state transitions. By doing so, we preserved the integrity of the temporal dynamics of the underlying data, ensuring that the total time for each state transition remained unchanged. This method allowed us to increase the number of training samples meaningfully, without artificially generating new state transitions or introducing redundancy.

We filtered training samples by location, model and forest state to reduce the bias resulting from multiple contributions from the same model and location to the simulation database. Specifically, the majority of simulations in the database are from two models (i.e. iLand and 4C), and some species were simulated more frequently than others (Grünig et al., 2024). To reduce the dominance of a single model and/ or species in the training dataset, we downsampled to a maximum of 100,000 training samples per location, while keeping as many different forest states as possible. Furthermore, we filtered out direct management signals from interventions such as thinnings or final harvests (e.g. by detecting and eliminating canopy height reduction by more than 2 m from the data) from the training samples, yet indirect management signals might still be included in the data. The resulting dataset comprised 2,750,456 million data points covering 5,445 distinct forest states. The number of training samples for the different species ranged from 34,759 (*Betula pendula*) to 250,634 (*Fagus sylvatica*; see Table S1).

*DNN architecture and cross-validations*

The implemented DNN was designed for multi-input, multi-output prediction tasks, specifically targeting forest state transitions and time to transition. The architecture integrates multiple data streams, including forest state, historical forest state (i.e. state before the previous transition), residence time, historical time (i.e. residence time before the previous transition), soil data, and climate inputs. The state and historical state inputs are first concatenated and processed through an embedding layer, followed by a flattening operation. This is succeeded by a dense layer with ReLU activation, reshaped to feed into a 1D convolutional layer, followed by max pooling and flattening. Historical state and time inputs undergo a masking process using a Lambda layer to apply predefined binary masks, simulating various levels of data availability during training and validation, as history of the states was not always available in the dataset. Further, climate data is processed through a series of dense layers (again with ReLU activation functions) and dropout regularization, then flattened. The processed inputs are concatenated and passed through a dense layer (ReLU activation), forming a combined representation. This combined input is further processed through a residual block, followed by branching into two separate pathways: one for predicting time to transition and another for predicting target states. The time to transition prediction branch includes a dense layer (ReLU activation), a residual block, dropout regularization and a final dense layer with Softmax activation over 10 target times (1 to 10 years). The state transition prediction branch consists of a dense layer (ReLU activation), a residual block, dropout regularization, and a final dense layer with softmax activation over 5,445 classes (i.e. all discrete forest states).

We tested the generalizability of the DNN with several cross-validation experiments. The first cross-validation was a random sample five-fold cross-validation. For this, we randomly split the training data into five equally sized chunks. We subsequently trained five DNNs, each time using four chunks for training and one chunk for validation. The second validation experiment was splitting the data with climate change scenarios (i.e. historical, RCP2.6, RCP4.5, RCP8.5). Four DNNs were trained, each using the data of three climate change scenarios for training and one for validation. We focused on the model selection as a third cross-validation experiment. We split the data by unique identifier code of the data in the harmonized forest simulation database. This code corresponds to the contributor of the data and in most cases refers to one model. Note that some contributors provided data from several models or had several unique identifier codes due to the large amount of data they contributed to the database. We randomly split to unique identifiers into five groups and again created five chunks from the data corresponding to those groups. The results of these cross-validation experiments showed that the DNN was generalizing well, although the variation between validation accuracies was higher for the climate scenario cross-validation and the model selection cross-validation.

Table S4: Cross validation with five equal sized chunks, randomly sampled. Results are shown for top prediction and top 2 predictions.

| CV | accurracy train state | accurracy train time | accurracy val state | accurracy val time |
|---|---|---|---|---|
| **1** | 0.8895 | 0.6591 | 0.8602 | 0.6024 |
| **2** | 0.8888 | 0.6576 | 0.8637 | 0.6075 |
| **3** | 0.8889 | 0.6603 | 0.8662 | 0.6099 |
| **4** | 0.889 | 0.657 | 0.8635 | 0.6113 |
| **5** | 0.8897 | 0.658 | 0.8646 | 0.6097 |
| **Mean** | 0.88918 | 0.6584 | 0.86364 | 0.60816 |
| **SD** | 0.0004 | 0.0012 | 0.002 | 0.0031 |

**Same for top 2**

| CV | accurracy train state | accurracy train time | accurracy val state | accurracy val time |
|---|---|---|---|---|
| **1** | 0.9681 | 0.8386 | 0.9519 | 0.7796 |
| **2** | 0.9681 | 0.8369 | 0.9529 | 0.7886 |
| **3** | 0.9682 | 0.839 | 0.9533 | 0.7924 |
| **4** | 0.9679 | 0.8373 | 0.9537 | 0.7899 |
| **5** | 0.9683 | 0.8378 | 0.9524 | 0.789 |
| **Mean** | 0.96812 | 0.83792 | 0.95284 | 0.7879 |
| **SD** | 0.0001 | 0.0008 | 0.0006 | 0.0044 |

Table S5: Cross validation with chunks split by climate change scenario. Chunk 1 is examples under historical climate conditions, chunk 2 RCP2.6, chunk 3 RCP4.5 and chunk 4 RCP8.5. Results are shown for top prediction and top 2 predictions.

| **ClimCV** | **accurracy train** | **accurracy train** | **accurracy val** | **accurracy val** |
|---|---|---|---|---|
| | **state** | **time** | **state** | **time** |
| **1** | 0.8846 | 0.6566 | 0.8598 | 0.5453 |
| **2** | 0.888 | 0.6578 | 0.7916 | 0.52 |
| **3** | 0.8941 | 0.6636 | 0.88 | 0.6354 |
| **4** | 0.9028 | 0.6714 | 0.8039 | 0.5406 |
| **Mean** | 0.892375 | 0.66235 | 0.83383 | 0.56033 |
| **SD** | 0.0069 | 0.0059 | 0.037 | 0.0444 |

| **Same for top 2** | | | | |
|---|---|---|---|---|
| **ClimCV** | **accurracy train** | **accurracy train** | **accurracy val** | **accurracy val** |
| | state | time | state | time |
| **1** | 0.9667 | 0.8351 | 0.9579 | 0.7404 |
| **2** | 0.9674 | 0.8369 | 0.9383 | 0.6808 |
| **3** | 0.9697 | 0.8438 | 0.9558 | 0.8206 |
| **4** | 0.9733 | 0.8516 | 0.9012 | 0.7203 |
| **Mean** | 0.969275 | 0.84185 | 0.9383 | 0.74053 |
| **SD** | 0.0026 | 0.0065 | 0.0227 | 0.051 |

Table S6: Cross validation with chunks unique identifier number. We randomly assigned the 38 unique identifiers to 5 groups and pooled data from those groups. Results are shown for top prediction and top 2 predictions.

| **CV_uniqueID** | **accurracy train** | **accurracy train** | **accurracy val** | **accurracy val** |
|---|---|---|---|---|
| | state | time | state | time |
| **1** | 0.9031 | 0.6707 | 0.6366 | 0.3807 |
| **2** | 0.9008 | 0.677 | 0.7726 | 0.4495 |
| **3** | 0.8863 | 0.6519 | 0.766 | 0.4856 |
| **4** | 0.8718 | 0.6447 | 0.9009 | 0.4973 |
| **5** | 0.8866 | 0.6562 | 0.6632 | 0.4113 |
| **Mean** | 0.88972 | 0.6601 | 0.7479 | 0.4449 |
| **SD** | 0.0114 | 0.012 | 0.0937 | 0.044 |

| **Same for top 2** | | | | |
|---|---|---|---|---|
| **CV_uniqueID** | **accurracy train** | **accurracy train** | **accurracy val** | **accurracy val** |
| | state | time | state | time |
| **1** | 0.9721 | 0.8524 | 0.7697 | 0.5286 |
| **2** | 0.9737 | 0.8551 | 0.8576 | 0.6323 |
| **3** | 0.9663 | 0.8336 | 0.8977 | 0.6705 |
| **4** | 0.9619 | 0.8194 | 0.9378 | 0.7797 |
| **5** | 0.9675 | 0.9358 | 0.7906 | 0.587 |
| **Mean** | 0.9683 | 0.85926 | 0.8507 | 0.6396 |
| **SD** | 0.0042 | 0.0404 | 0.0633 | 0.0845 |

As a fourth validation experiment, we focused on class imbalance in the training data, and tested the ability of the DNN to generalize between states of high and low frequency. Our dataset consists of 5,445 different forests states, of which the majority occurs less than 100 times and less than 10% occur more than 1,000 times. We therefore tested how the DNN performs on five different classes of state frequency (1 – 19, 20 – 99, 100 – 499, 500 – 999, >= 1000). For this, we randomly split the training data into five equally sized chunks and subsequently trained five DNNs, each time using four chunks for training and one chunk for testing. We then run 20 iterations of sampling 100,000 training samples, each time subsetting the state frequency classes. We then predicted state transitions for each frequency class of the test data and obtained the accuracy. While the DNN performed best on the most frequent states, the accuracy for the rarest states was 0.81 compared to the overall accuracy of 0.867 (Figure S19).

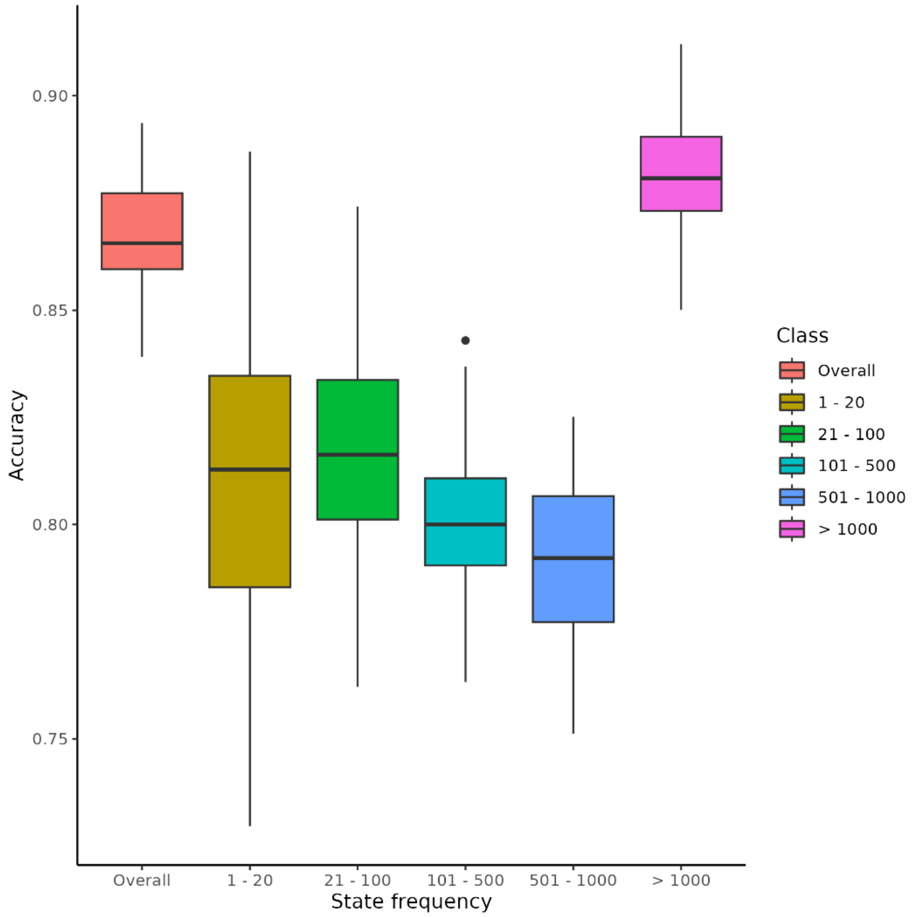


*Figure S19: DNN performance on five different classes of state frequency. A DNN was trained on 80% of the data and tested with 20%. From the test data, repeated sampling (10 times repetition) of data was used to obtain training samples to predict state transitions. Training samples were filtered by the frequency class and accuracy was measured as performance metric per class and overall.*

We further used an approach of explainable AI to assess the variable importance for DNN predictions. We applied variable permutation for all groups of variables, forest state, forest state history, residence time, residence time history, soil variables and climate variables. We trained a DNN on 80% of the data and run the permutation analysis on 20% random holdout data. Each variable group was tested by shuffling the data of the focal variable before predicting state transitions and measuring prediction performance. We found that the current forest state was the most important and the history of the forest state (i.e. the last three forest states) the second most important predictor. This aligns with the expectations, as for predicting future forest states their current and past states are strong indicators of their likely future states. Further the information on current and past state embeds the legacy of the past and provides information whether the forest conditions are stable over time or changing rapidly. Climate was the third most important variable, showing that the DNN predictions are climate sensitive. Residence time and the residence time history showed a slightly lower importance than climate and the soil variables showed the lowest contribution to the prediction performance.

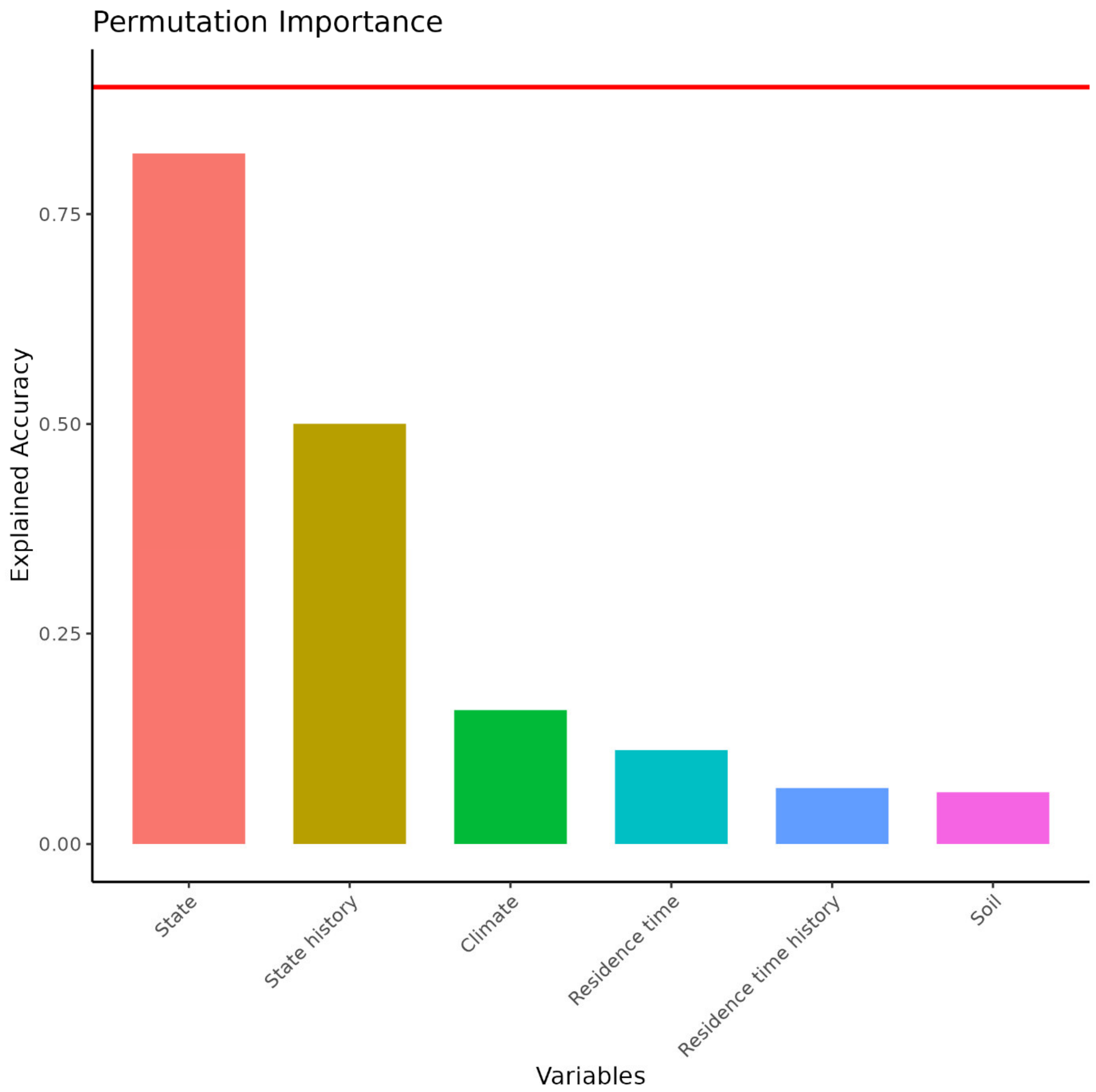


*Figure S20: Permutation importance on holdout data (20% randomly sampled) for the six variable groups. The red line shows the overall accuracy.*

*Uncertainty estimates*

All DNN predictions were probabilistic, i.e. probabilities for the predicted future forest state were obtained from the DNN, and the most probable state analyzed further in the context of our research questions (see above). To further elucidate uncertainties of our DNN meta-model, we examined the probability of the predicted forest state transitions and times to transition across the range of each species (Fig. S13 – S16). These maps indicate the degree of certainty that the DNN has in a projected state transition for a given cell. They highlight that for species where high amounts of training data were available (i.e. European beech, Norway spruce, Scots pine) the DNN tended to be more confident than for species with less training data. The spatial variation in the thus obtained uncertainty maps suggests that regions poorly covered by underlying simulation data had higher uncertainty in DNN predictions. For instance, predictions for silver birch in Eastern Europe had lower confidence due to the limited availability of training data for that species in this region.

*Climate data*

We downloaded climate data from three different global circulation models (GCMs; MPI-M-MPI-ESM-LR (Brovkin et al., 2019), ICHEC-EC-EARTH (EC-Earth Consortium, 2014), NCC-NorESM1-M (NORCE Norwegian Research Centre & Norwegian Meteorological Institute, 2011), all downscaled with the SMHI-RCA4 RCM (Strandberg et al., 2014)). To select GCMs, we filtered all available GCMs for those which provided consistent data (i.e. daily data for the needed variables and data for all years from 1981 to 2100) for all three RCPs. Further, we based the GCM selection on model skill evaluation and gradient coverage of temperature and precipitation conditions based on the GCMeval tool (Parding et al., 2020). The selected GCMs represent rather conservative change in temperature and precipitation (Figure S21.

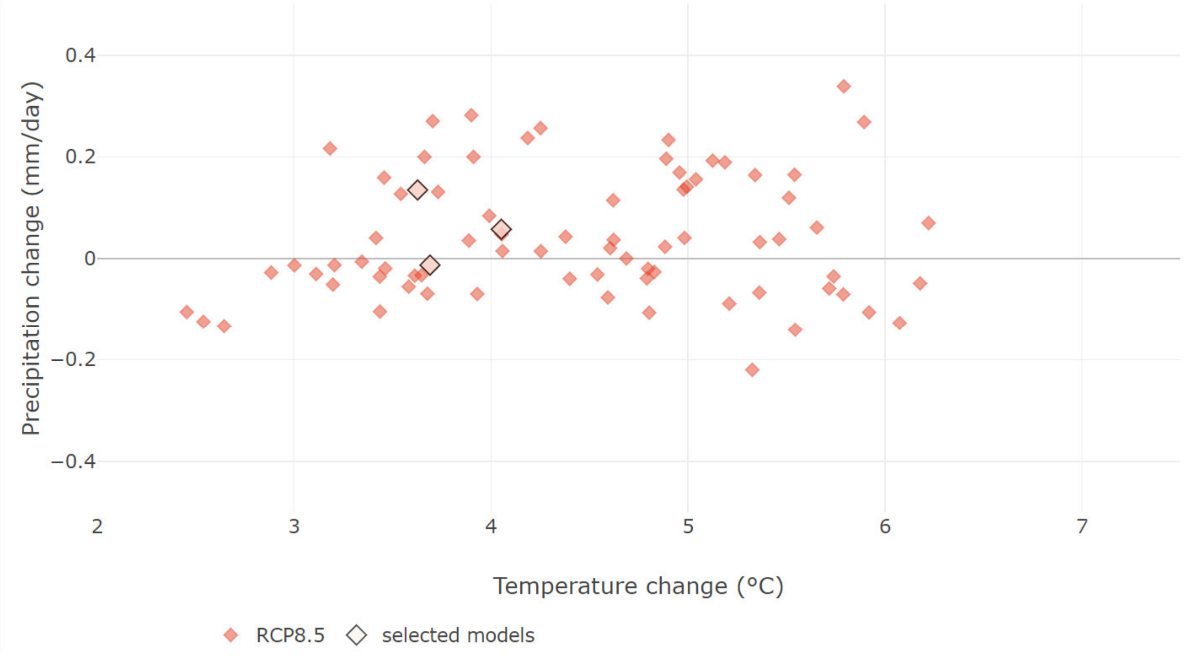


*Figure S21: GCM selection based on the GCMeval tool (Parding et al., 2020). Red points show all CMIP5 models on gradients of precipitation change and temperature change under the RCP8.5 scenario. The three selected models are highlighted. While the selected models are not covering the full gradient in temperature and precipitation contained within all* EURO-CORDEX simulations*, they account for some variation and are rather conservative with regard to the changes in both variables.*

*Climate compressor approach*

We run 3PG simulations using iLand (Seidl et al., 2012) to estimate Net Primary Productivity (NPP) for ten tree species (*Picea abies, Abies alba, Larix decidua, Pinus sylvestris, Fagus sylvatica, Quercus robur, Betula pendula, Quercus ilex, Pinus pinaster, Pinus halepensis*) in Europe based on daily climate and soil conditions. For this, we stratified the climate (using the previously described historical EURO-CORDEX data) with annual precipitation, mean annual temperature and temperature seasonality into 207 distinct climate bins. From each climate bin, we sampled up to 10 datapoints (i.e. when less datapoints in a bin where available we obtain all), resulting in 1360 climate datapoints. We further stratified soil conditions across Europe based on pan-European soil data on soil texture, water holding capacity and plant-available nitrogen (described in the Methods section Soil conditions), resulting in 56 soil bins. In total this resulted in 76,160 combinations of climate and soil conditions, for which we ran NPP simulations for 120 years under three different GCMs and three RCP Scenarios. With the outputs of the NPP simulations, we trained a DNN to predict NPP based on the daily climate data and soil data. The network consisted of two input branches: one for daily climate data across a year (365 days, 4 features) and another for environmental site variables (5 features). The climate branch aimed at reducing the dimensionality of the climate data towards a small number of units in the last dense layer before the output (Fig. S22). The environmental branch directly fed into the network. Both branches were concatenated and passed through a series of dense layers with using GELU activation functions. The output layer consisted of 10 linear units to predict NPP for each tree species. The model was compiled with the Adam optimizer (learning rate 0.001) and mean squared error as the loss and evaluation metric. We tested different numbers of units in the last dense layer of the climate branch and found that the DNN performed best using 14 units. The DNN was trained on 50% of the data and 33% were used for validation and 17% for testing. The DNN performed well with an MSE of 0.0137 on the test dataset. We used the trained DNN to predict NPP from daily climate data and at the same time extracted the reduced climate data (i.e. climate indices) from the dense layer at the end of the climate branch. The 14 climate indices and the 10 NPP values were used as climate data in the main DNN.

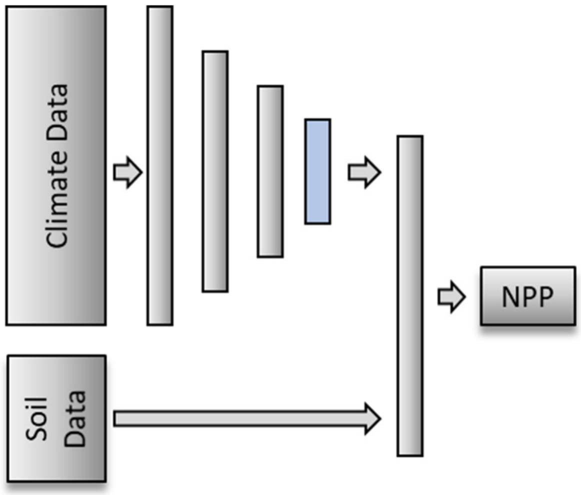


*Figure S22: Schematic display of the climate compressor DNN. The inputs contain climate and soil data. Climate data was reduced in dimensionality by passing through several layers until a dense layer with 14 units. This dense layer was used to extract the climate indices.*
